# Respiratory synchrony and individual differences causally influence dyadic interpersonal coordination

**DOI:** 10.1101/2025.05.02.651958

**Authors:** Wenbo Yi, Caroline Palmer

## Abstract

Human interpersonal coordination can yield synchronization at multiple timescales, including behavioral (auditory-motor) and physiological (respiratory and cardiac) levels; yet the causal relationship among these levels is poorly understood. By comparing dyadic melody perception and production, we demonstrate that physiological synchrony is not merely a byproduct of shared perception, as it increases significantly during joint production relative to joint perception or to silence. Perturbing dyads’ behavior or respiration revealed distinct causal effects: respiratory perturbations impaired both dyadic respiratory and behavioral synchrony, whereas auditory-motor perturbations disrupted only dyadic behavioral synchrony. Individual differences further shaped synchrony: partners who shared similar spontaneous rates achieved better behavioral synchrony, and partners with more similar resting heart rates exhibited stronger cardiac synchrony in joint production. These findings disentangle the relationships among levels of human synchrony, reveal directional entrainment processes between respiratory and behavioral synchrony, and highlight the pivotal role of individual differences in interpersonal coordination.

## 1. INTRODUCTION

Human interpersonal coordination often incorporates both intentional and spontaneous (unintentional) synchronization of individuals’ actions. Synchrony is considered to be a fundamental, evolutionarily based mechanism that facilitates social cohesion and bonding among people^1–6^. In addition to behavioral synchrony, physiological synchrony between individuals, such as alignments in their respiratory and cardiac patterns, can coincide with group cohesion and shared intentions^7–16^. Increased physiological synchrony has been observed among members of musical groups, including choral singers^17^, string quartet musicians^18^, and drumming groups^10,19^, when music performance is compared with a silent baseline. Despite these findings, causal relationships among behavioral and physiological synchrony that arise in interpersonal interactions among individuals are not well-understood.

Studies of interpersonal coordination often assume that respiratory or cardiac alignment between individuals emerges when they perform a similar task together; how behavioral synchrony facilitates that physiological synchrony is unclear^20,21^. Some research suggests that stronger cardiac or respiratory synchrony occurs between individuals who perform a behavioral synchronization task together, compared with a baseline condition^17,19,22^. However, these findings did not establish a causal link between the behavioral and physiological variables, as the directional influence of one variable on the other was not manipulated^23,24^. One study directly examined the causal role of auditory-motor synchrony in cardiac synchrony by comparing drummer trios who synchronized their beats with either predictable or unpredictable acoustic cues. Drummers exhibited greater behavioral asynchronies when presented with unpredictable auditory cues, but their cardiac synchrony remained unchanged across the acoustic conditions^10^. These mixed findings raise a critical question: Whether physiological synchrony that sometimes arises between individuals is a byproduct of their shared perceptual experience (i.e., participants are experiencing similar events in shared environments^20,25^) or is instead the result of direct behavioral interactions between participants.

Another unsolved issue is whether physiological synchrony is necessary to facilitate interpersonal coordination. It is well-established that physiological rhythms can modulate human cognitive processes^26–28^. For example, different phases of respiratory activity can impact behavior: the inhalation phase has been shown to enhance memory retrieval^29^, visuospatial accuracy^30^, and auditory discrimination abilities^31^ relative to the exhalation phase. As well, the cardiac systole phase (heart muscle contraction) is often accompanied by baroreceptor noise that can inhibit visual and auditory perception while simultaneously enhancing motor excitability^26^. Conversely, the diastole phase (heart muscle relaxation) is associated with reduced motor activity but enhanced sensory evaluation^26,27,32,33^. These findings raise important questions about whether and how physiological rhythms influence joint behavior among individuals. The current study manipulated groups’ respiratory synchrony as an independent variable to test its causal influence on groups’ behavioral synchrony, and vice versa.

Both behavioral and physiological rhythms show large and consistent differences across individuals^21,34–38^. For example, humans tend to produce reliable and characteristic differences in movement rates when talking, walking, performing music, or singing^20^. These optimal movement frequencies, shown to require the least energy expenditure, have been modeled as the natural frequencies of oscillators that represent preferred coordination states within a system, known as the Spontaneous Production Rate (SPR)^35,39–42^. Partners with similar SPRs perform more synchronously in joint tasks^36^, a finding that aligns with the predictions of coupled oscillators with different natural frequencies; during entrainment in joint actions, oscillators with similar frequencies will synchronize more^43^. Similarly, individual differences in resting heart rates have been observed that remain consistent over time^44^, but can differ across individuals by as much as 70 beats per minute. If a dynamic entrainment process occurs between oscillations with different natural frequencies, then dyad members with similar physiological baselines (such as resting heart rates) may synchronize more than those with different physiological baselines.

We investigated causal relationships between behavioral and physiological synchrony in two-person productions of melodies, while unpredictable perturbations disturbed their auditory-motor and respiratory rhythms. Amateur musicians were randomly paired to perform a musical synchronization task (Figure 1A). Both respiratory and cardiac rhythms were measured, as individual respiratory rhythms are known to be coupled with cardiac rhythms in a range of tasks, including rest, sleep, and physical activity^45–47^. Participants first completed a standardized individual melody production task^35,36,39–41^ to quantify their SPRs as a measure of behavioral individual differences. Next, participants sat quietly together for a silent Baseline measurement of their resting heart rates and breathing rates. Then participants completed a Perception task, during which they heard the same auditory sequences they would later produce. Finally, participants performed the Production task, beginning with the Normal condition, in which they synchronized their melody production with their partner (Figure 1B). Sounded perturbations occurred at unpredictable stimulus locations in two conditions: In the Auditory condition, a sounded cue signaled that partners should restart the melody production immediately. In the Respiratory condition, the same sounded cue signaled that partners should take a rapid deep breath, thereby resetting their respiration rhythms. A final Normal condition was collected last.

**Figure 1.**
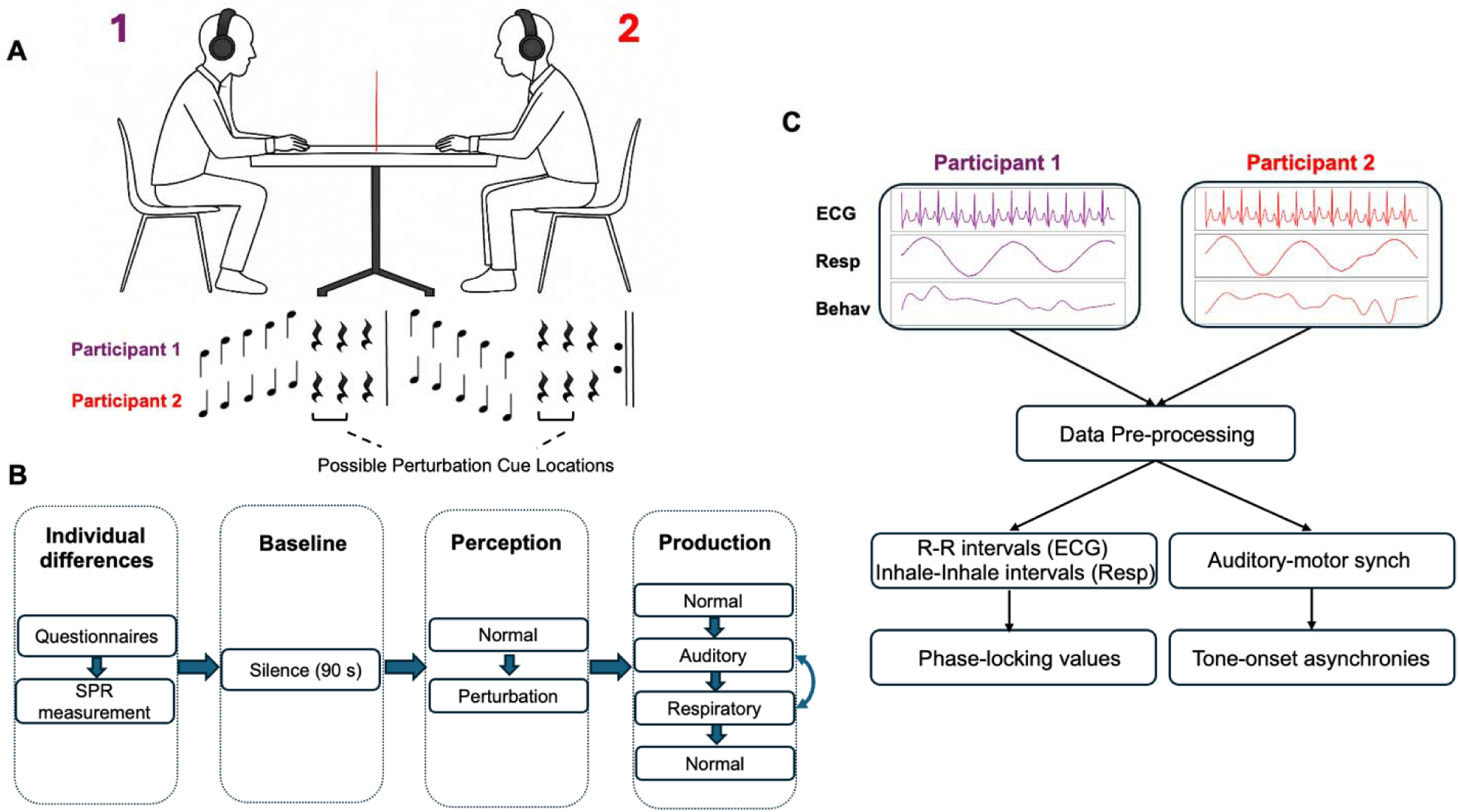
Experimental paradigm. **A)** Two participants sat face-to-face with a visual barrier occluding finger movements. Each participant produced an auditory sequence, with possible perturbation cue locations indicated. **B)** Overview of the experimental procedure, including individual tasks (questionnaires, spontaneous production rate measurement), baseline task while seated together (90 s of silence), perception tasks while seated together (listening to the Normal melody without cues and the Perturbation melody with cues), and production tasks. Production conditions included: *Normal* (melody without cues), *Auditory* (melody with auditory cues prompting participants to reset to the start), and *Respiratory* (melody with auditory cues prompting participants to inhale deeply and quickly to reset respiration). The Normal condition was repeated, and the order of Auditory and Respiratory conditions was counterbalanced across dyads. **C)** Data analysis pipeline. Physiological (ECG and respiration) and Behavioral (tone onset) data were recorded from each participant. Data pre-processing procedures were applied to ECG, respiratory, and behavioral signals (see Methods for details).

We predicted that the Production task, in which both partners contributed to the shared synchronization goal, would yield higher levels of respiratory/cardiac synchrony than the Baseline and Perception tasks. The Normal Production condition (with no perturbations) was expected to yield the greatest behavioral and respiratory/cardiac synchrony. The Auditory Production condition (auditory-motor perturbations) was anticipated to disrupt behavioral synchrony and to cause reduced respiratory/cardiac synchrony between partners. Finally, the Respiratory Production condition (respiratory perturbations) was expected to reduce both behavioral and respiratory synchrony, compared with the Normal and Auditory conditions.

## 2. RESULTS

### 2.1 Auditory-motor synchronization in tone onsets

#### 2.1.1 Perturbation effects on auditory-motor synchronization

Dyadic behavioral synchronization was measured by the absolute asynchronies between the partners’ tone onsets (intended as simultaneous). There were no significant differences in absolute asynchronies between the first and last Normal conditions (*t*(30) = -0.003, *p* = 0.998), and the two conditions were combined. The repeated measures ANOVA on the absolute asynchronies by Task revealed a significant main effect (Figure 2A, B), *F*(2,60) = 14.08, *p* < .001, *η²* = 0.319. Post-hoc comparisons showed that absolute asynchronies in the Normal condition were significantly smaller than those in the Auditory condition, *t*(30) = -2.505, *p_holm_* = 0.018, *d* = -0.274, and the Respiratory condition, *t*(30)= -4.602, *p_holm_*< .001, *d* = -0.870. Additionally, asynchronies in the Auditory condition were significantly smaller than those in the Respiratory condition, *t*(30) = -3.113, *p_holm_* = .008, *d* = -0.596. Thus, participants synchronized their auditory-motor behaviors highest in the Normal condition, followed by the Auditory condition, with the lowest synchronization in the Respiratory condition. The mean inter-tap intervals showed that participants produced the melodies significantly slower in the respiratory condition than in the Normal and Auditory conditions (See SI.1).

**Figure 2.**
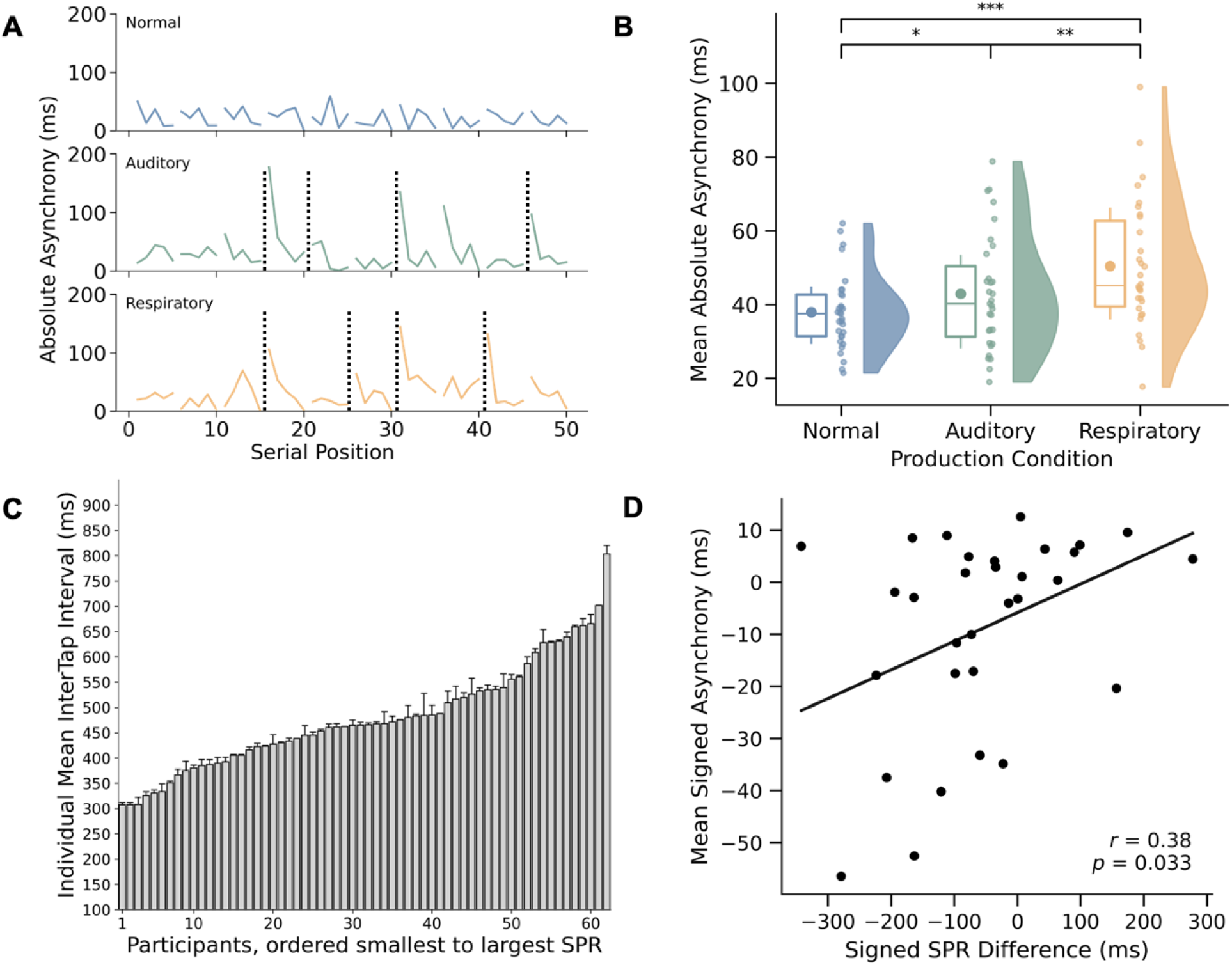
Behavioral measures from Spontaneous Production Rate and Production task. **A)** Example trials from one dyad showing auditory-motor synchronization across Production conditions (Normal, Auditory, Respiratory). Absolute asynchronies were computed as the absolute difference between tone onsets from Participant 1 and Participant 2. Dashed lines indicate perturbation cue locations; gaps denote silent beats (rests) within a trial. **B)** Mean absolute asynchronies across dyads in the three Production conditions. Box plots display medians (horizontal lines) and means (circles in boxes). Significance markers: *p* < 0.05 (\**), p < 0.01 (**\*\****), *p* < 0.001 (***). **C)** Mean intertap intervals (ms) from the Spontaneous Production Rate (SPR) task, ranked from fastest to slowest across participants. Error bars indicate the standard error of the mean. **D)** Correlation between mean signed asynchrony (Participant 1’s higher-pitched tone onsets - Participant 2’s lower-pitched tone onsets, in ms) and the dyad’s SPR difference (Participant 1 - Participant 2, in ms). A significant positive correlation was observed (Pearson *r* = 0.38, *p* = 0.033); the regression line is shown.

To determine whether the perturbation effects were short-lived (i.e., only affecting tone synchronization immediately after the perturbation cue), we removed the first and second asynchronies following each cued location from the data. The same significant findings were obtained, with asynchronies following the same pattern (See SI.2). Together, these findings suggest that both auditory and respiration perturbations negatively impacted partners’ behavioral synchronization, with respiratory disruptions yielding the largest reduction.

#### 2.1.2 Individual differences in auditory-motor synchronization

We examined the dyads’ asynchronies in terms of each partner’s Spontaneous Production Rates (SPR) observed while producing a familiar melody, to quantify individual differences. Each participant’s mean inter-tap interval (ITI) per trial from the SPR Measurement task demonstrated consistent inter-individual rate differences (Figure 2C). We then tested whether the dyad partners’ differences in spontaneous production rate (SPR) predicted their dyadic behavioral synchronization Following established procedures in previous studies^39,41,48^, we calculated signed asynchronies in tone onsets within each dyad (Participant 1 - Participant 2) and correlated them with the signed SPR differences between dyad members (Participant 1 - Participant 2) in the Normal (nonperturbed) condition. The partners’ signed asynchronies correlated positively with their signed SPR differences, Pearson’s *r* = 0.38, *p* = 0.03 (Figure 2D). The dyad partner with the faster spontaneous rate tended to precede in tone asynchronies the partner with the slower spontaneous rate. Consistent with previous results^41,48^, the larger the partners’ differences in spontaneous rates, the larger their tone asynchronies were in the direction predicted by the SPR values.

### 2.2 Perturbation effects on physiological synchronization

Respiratory and cardiac synchrony between the two dyad members was quantified using Phase Locking Values (PLV), which range from 0 to 1, with 1 indicating perfect phase synchronization^49,50^. Respiratory and cardiac PLVs showed no significant differences between the first and last Normal conditions (see SI.3) or between the Perception-Normal (without perturbation cue) and Perception-Perturbation (with perturbation cues) conditions (see SI.4). Therefore, we averaged the two Normal conditions and the two Perception conditions into a single Normal and Perception task, respectively. The Production task conditions (including Normal, Auditory, and Respiratory conditions) were contrasted with the Baseline and Perception tasks to examine physiological differences across the tasks. Finally, we analyzed differences among the Normal, Auditory, and Respiratory conditions within the Production task to assess the effects of perturbations on physiological activities.

#### 2.2.1 Respiratory synchronization

We assessed respiratory synchrony between dyad members using Phase Locking Value (PLV) computed on normalized respiration amplitudes by Task. There were significant differences among the Baseline, Perception, and Production tasks, *F*(2,60) = 3.251, *p* = 0.046, *η²* = 0.098. Linear contrasts were applied to test the hypothesized lower Baseline PLV values than in the other conditions; as expected, the Baseline PLV was significantly lower than the Production PLV, (*t*(30) = -2.144, *p* = 0.04, *d* = -0.385). The difference between Baseline and Perception PLVs was not significant (*t*(30) = -1.658, *p* = 0.108). There were no significant differences between the Perception and Production conditions (*t*(30) = 1.060, *p* = 0.30).

We next analyzed the PLV values in the three Production conditions (Normal, Auditory, Respiratory), which also indicated a significant main effect, *F*(2,60) = 18.619, *p_holm_* < .001, *η²* = 0.383. As expected, the Respiratory condition had the lowest PLV compared to the Normal (*t*(30) = 5.183, *p_holm_* < .001, *d* = 0.955) and Auditory (*t*(30) = 4.53, *p_holm_* < .001, *d* = 1.00) conditions (Figure 3A). There was no significant difference between the Normal and Auditory conditions (*t*(30) = -0.318, *p_holm_* = 0.753). Surrogate analyses demonstrated that the observed dyads showed significantly higher PLV compared to randomly reassigned dyads (*p* < 0.0042) or to randomly reassigned conditions (*p* < 0.001, see SI.5), confirming that respiratory synchrony emerged in interpersonal coordination and was manipulated by the specific conditions (Normal, Auditory, Respiratory).

**Figure 3.**
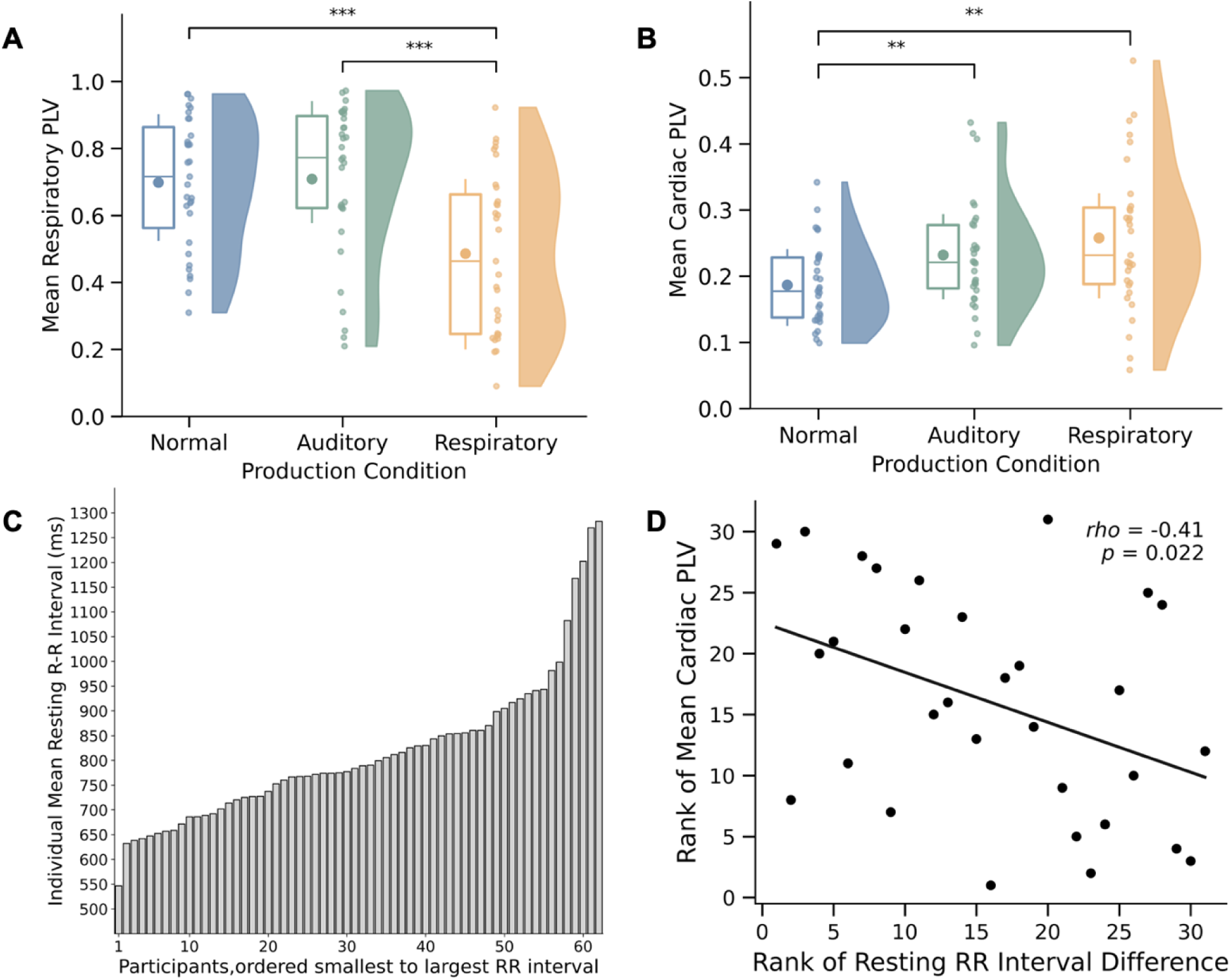
Respiratory and Cardiac measures from Production task. **A)** Mean respiratory phase-locking values (PLVs) between dyad members across Production tasks (Normal, Auditory, Respiratory). Box plots show medians (horizontal lines) and means (circles in boxes). Significance markers: *p* < 0.05 (\**), p < 0.01 (**\*\****), *p* < 0.001 (***). **B)** Mean cardiac PLVs across Production tasks (Normal, Auditory, Respiratory). Box plots indicate medians (horizontal lines) and means (circles in boxes), significance markers: *p* < 0.05 (\**), p < 0.01 (**\*\****), *p* < 0.001 (***). **C)** Individual mean resting heart rates (mean R–R interval, in ms) recorded during the Baseline task, ranked from fastest (shortest interval) to slowest rate (longest interval). **D)** Correlation between absolute differences in the ranks of resting heart rate between dyad members (Participant 1 – Participant 2 in R–R interval, ms) and the rank of their mean cardiac PLV in the Normal condition. A significant negative correlation was observed (Spearman’s rho = –0.41, *p* = 0.022); the regression line is shown.

In sum, these findings confirm that the respiratory perturbation manipulation significantly disrupted respiratory synchronization in interpersonal coordination. The non-significant respiration PLV differences between the Normal and Auditory conditions suggest that perturbing the dyads’ behavioral synchronization did not affect their respiratory synchronization.

#### 2.2.2 Cardiac synchronization

Phase Locking Values (PLV) on heart rates were analyzed next to measure cardiac synchronization between dyad members. The repeated-measures ANOVA by task indicated significant differences in cardiac PLVs (*F*(2, 60) = 3.509, *p* = 0.036, *η²* = 0.105). The PLVs were highest in the Production task compared to the Perception task (*t*(30) = -2.471, *p* = 0.01, *d* = -0.483) and Baseline task (*t*(30) = -2.737, *p* = 0.01, *d* = -0.585). There was no significant difference between the Perception and Baseline tasks (*t*(30) = -0.354, *p* = 0.725). Thus, joint melody production enhanced cardiac synchronization, and this effect was not driven by perception alone, as Perception and Baseline PLVs did not differ.

The same analysis was performed on the PLV values from the Normal, Auditory, and Respiratory conditions. Results showed significant differences between the three Production conditions, *F*(2, 60) = 7.688, *p* = 0.001, *η²* = 0.204 (Figure 3B). The PLVs in the Normal condition were significantly lower than in both the Auditory (*t*(30) = -3.166, *p_holm_* = 0.007, *d* = -0.52) and Respiratory (*t*(30) = -3.712, *p_holm_* = 0.003, *d* = -0.816) conditions. There was no significant difference between Auditory and Respiratory perturbation conditions (*t*(30) = -1.230, *p_holm_* = 0.228). Surrogate and Monte-Carlo analyses showed that the observed dyads’ cardiac synchronies were significantly higher than randomly assigned dyads (*p = 0.019*) or re-assigned conditions (*p* < 0.001), suggesting that the observed synchrony was not attributable to chance or solely depend on shared responses to external stimuli, but instead reflected genuine interpersonal coordination shaped by specific dyadic interactions or task context. (See SI.5 for details).

#### 2.2.3 Individual differences in cardiac synchronization

To examine whether individual differences influence cardiac synchrony, we assessed the correlation between the absolute difference in partners’ resting heart rates measured during Baseline and their PLVs in the Normal synchronization condition. Similar to previous findings^44^, the mean resting heart rates (in R-R intervals) showed large but consistent inter-individual differences (Figure 3C). A Spearman correlation was conducted (the Shapiro-Wilk test indicated violation of the normality assumption); the absolute difference in partners’ resting heart rates (measured as mean R-R interval differences) was negatively correlated with the dyad’s cardiac PLV in the Production task (Spearman *rho* = -0.41, *p* = 0.022, Figure 3D). Thus, greater similarity in dyad members’ resting heart rates was associated with stronger cardiac synchrony during Production. The correlations between the same measures in the Baseline and Perception tasks were not significant, suggesting that the partners’ joint actions, rather than shared perception facilitated cardiac entrainment between individuals (see SI.6 for details).

In sum, the dyad’s joint action facilitated respiratory and cardiac synchrony^10,17,18^, and perturbations demonstrated that resetting either behavioral or respiratory synchronization promoted cardiac synchrony. These findings suggest that dyadic cardiac synchronization can be independent of both auditory-motor and respiratory synchronization.

### 2.3 Correspondences between dyadic behavioral, respiratory, and cardiac synchrony

We tested correspondences between the behavioral and physiological synchronization measures in the Production task. Dyadic auditory-motor asynchronies were negatively correlated with dyadic respiratory PLV measures in the Normal condition (Pearson *r* = -0.48, *p* = 0.006), indicating that dyads with greater respiratory phase-locking showed higher behavioral synchronies. Importantly, this relationship persisted even when respiratory synchronization was perturbed (Respiratory condition: Pearson *r* = -0.50, *p* = 0.005, Figure 4A), but disappeared when behavioral synchronization was perturbed (Auditory condition: Pearson *r* = -0.161, *p* = 0.387). These findings further support the previous main effect that disrupting respiration impacts behavioral synchrony, whereas disrupting behavioral synchrony does not significantly alter respiration. When dyads breathed in synchrony, they were more likely to align their behavior more precisely.

**Figure 4.**
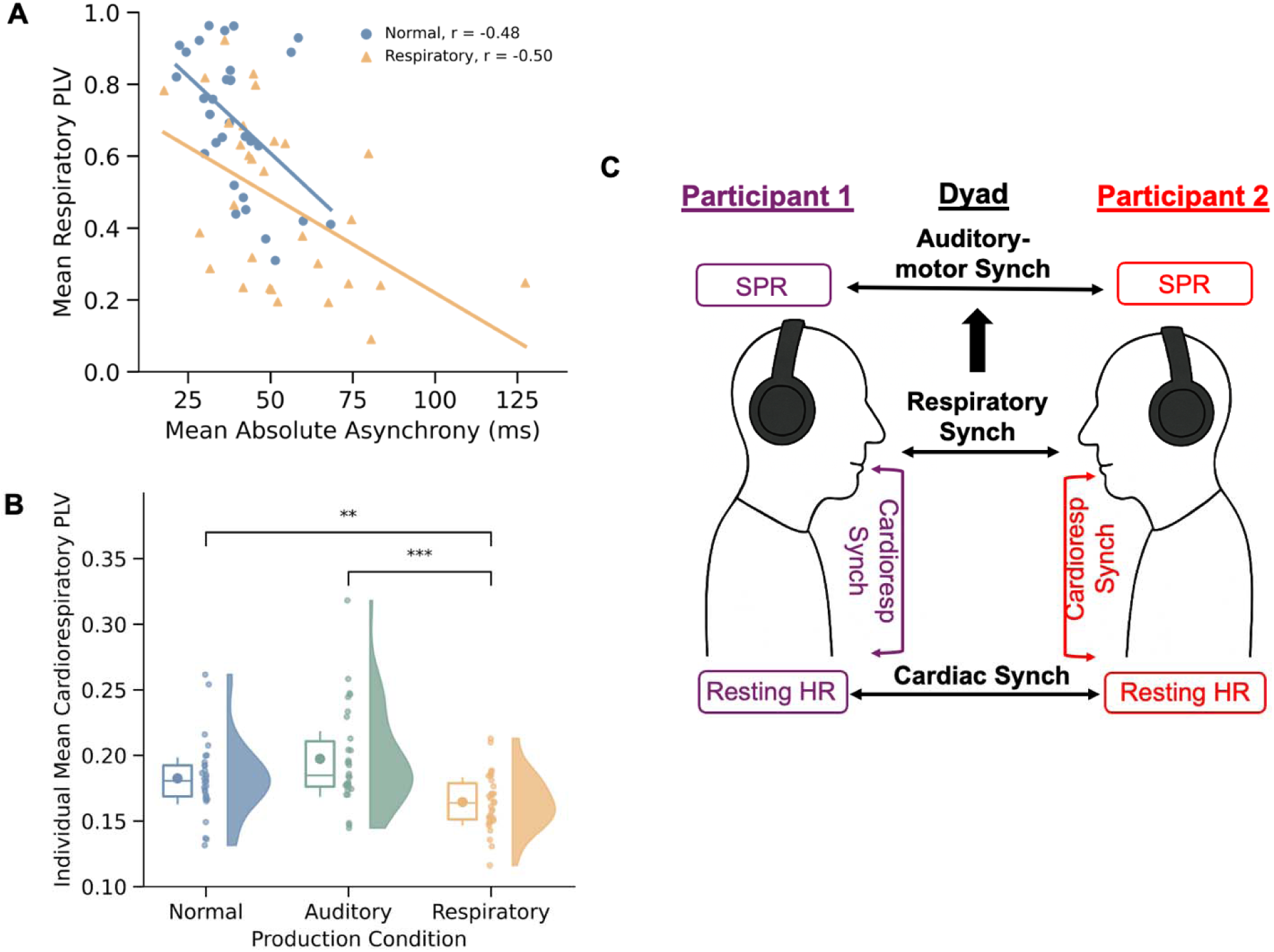
Dyadic Respiratory and Individual Cardiorespiratory measures in Production task. **B)** Negative correlations between participants’ mean absolute asynchronies and their respiratory PLVs in both the Normal (Pearson *r* = –0.48, *p* = 0.006) and Respiratory (Pearson *r* = –0.50, *p* = 0.005) conditions, indicating a tight link between respiratory synchrony and coordinated behavior. **C)** Mean cardiorespiratory PLVs within individuals across Production conditions. Box plots show medians (horizontal lines) and means (circles in boxes). Significance markers: *p* < 0.05 (\**), p < 0.01 (**\*\****), *p* < 0.001 (***). **D)** Schematic summary of observed relationships between auditory–motor, cardiac, and respiratory Dyadic synchrony (middle, black), and individual differences in rates and cardiorespiratory measures. Solid arrows indicate observed directional influences.

Dyadic tone onset asynchronies did not correlate significantly with the dyads’ cardiac PLV in the Normal synchronization condition or the Auditory or Respiratory (perturbation) conditions (see SI.7). In addition, no significant correlations were observed between the dyads’ cardiac and respiratory PLVs in any of the Production tasks (see SI.8).

### 2.4 Individual physiological rhythms and cardiorespiratory coupling

To investigate whether cardiac and respiratory coupling were linked within individuals in the Production task. Considering the large frequency difference between cardiac and respiratory signals, we quantified individuals’ cardiac–respiratory phase synchrony using the n:m phase-locking value, where m was fixed at 1, and n was determined by rounding the ratio of cardiac to respiratory frequency to the nearest integer for every trial^45,47,50^ (see Methods). Repeated-measures ANOVA on the PLVs revealed a significant effect of Production condition (*F*(2, 60) = 14.18, *p* < 0.001, *η²* = 0.321, Figure 4B). Post hoc comparisons showed that individuals’ cardiorespiratory PLVs were significantly lower in the Respiratory condition compared to both the Normal condition (*t*(30) = 3.35, *p_holm_*= 0.004, *d* = 0.623) and the Auditory condition (*t*(30) = 5.50, *p_holm_* < 0.001, *d* = 1.031). No significant difference was observed between the Normal and Auditory conditions (*t*(30) = –1.94, *p_holm_* = 0.062). To confirm that individual cardiorespiratory PLVs reflected genuine within-individual coupling, we conducted a surrogate analysis in which each participant’s cardiac time series was reassigned to the respiration time series of another individual within each Production condition. A Monte Carlo simulation revealed that PLVs were significantly higher in the observed individuals’ cardiorespiratory pairings compared to the surrogate pairings (*p* < 0.001; see SI.9 for details), indicating reliable phase coupling within individuals’ cardiorespiratory systems.

In sum, individual-level physiological dynamics indicated that respiratory perturbations led to a decoupling of cardiac and respiratory rhythms. In contrast, auditory-motor perturbations did not significantly alter individual physiological dynamics.

## 3. Discussion

Human interpersonal synchronization occurs at multiple levels; the present study reported a directional relationship between behavioral and physiological synchronization in dyads while they performed a joint task in a novel perturbation paradigm. We replicated previous findings^17–19,51,52^ that joint group activity promotes dyadic cardiac and respiratory synchronization compared to a quiet baseline. Critically, the dyads’ increased physiological synchronization was not simply due to shared perceptual experiences; no significant change or causal relationship was observed in the Perception task, which required the dyads to perceive and respond to the same auditory stimuli as they heard during joint production. Furthermore, we reported the first causal evidence linking respiratory synchrony to auditory-motor synchrony; when dyads breathed more synchronously, they synchronized their auditory-motor behavior more accurately. The study also provided evidence that the dyads’ individual differences influenced interpersonal synchronization at physiological levels. Dyads with smaller individual differences in Spontaneous Production Rate achieved better behavioral synchrony in the Production tasks^40,41,48^. Similarly, dyads with smaller differences in resting heart rate exhibited higher cardiac synchrony in joint production. Together, these findings provide new insights into the dynamic entrainment processes underlying person-to-person synchronization at both behavioral and physiological levels, which we review next.

First, we demonstrated a causal, unidirectional influence of respiratory synchrony on auditory–motor synchrony (Fig. 4C). Disruptions to respiratory synchrony during the Respiratory condition significantly reduced the dyads’ auditory–motor synchronization, whereas disruptions to auditory–motor synchrony during the Auditory condition had no measurable effect on dyads’ respiratory synchrony. This asymmetry supports a directional influence from respiratory to behavioral synchrony. Follow-up correlation analyses further indicated the critical role of respiratory synchrony in interpersonal coordination: a significant negative correlation between dyads’ respiratory PLVs and auditory-motor asynchronies was observed in the Normal condition, and this relationship persisted in the Respiratory condition, despite direct perturbation of breathing patterns. These findings reveal a millisecond-level correspondence between temporal behavioral and physiological coordination, extrapolating beyond prior evidence that compared synchrony in joint action with baseline conditions only^17,18^.

Second, heightened cardiac synchrony between dyad members was observed in all perturbation conditions, despite the dyads’ increased auditory-motor asynchronies in those conditions. Cardiac synchrony appeared to be independent of the dyads’ auditory-motor synchrony, similar to previous studies of group drumming^10^. Instead, the elevated cardiac synchrony may have been driven by the resetting demands imposed by the perturbation cues. We propose that the perturbation cues may have modulated the dyads’ dynamic attending processes, leading to shared fluctuations in attention and arousal throughout the trial^53^. This interpretation is supported by findings that group cardiac synchrony decreases in distracted-listening conditions compared to attentive-listening conditions during audiovisual narratives^54^. Although increased cardiac synchrony is often interpreted as an index of shared positive affect and group cohesion^7,9,10,15,55^, our findings suggest that cardiac synchrony can also reflect negative outcomes, such as the dyads’ increased behavioral asynchronies. Research findings on mother-infant interactions indicate that cardiac synchrony can arise during shared stress responses or stressful social interactions^56^. Rather than an index of shared positive affective valence among group members, cardiac synchrony is more likely a general physiological marker of shared attentional fluctuations.

Previous findings have focused on individual cardiorespiratory coupling and how that coupling changes in individual psychophysiological states and tasks^45,46,57^. In the current study, individuals’ cardiorespiratory phase coupling was reliably maintained in the Normal and Auditory conditions but significantly reduced under Respiratory perturbations, reflecting some cardiorespiratory dissociation when breathing conditions were altered. This within-individual reduction coincided with decreased dyadic respiratory synchrony and elevated dyadic cardiac synchrony relative to the Normal condition. Consistent with previous findings, this demonstrates that dyadic heart rate synchronization is always not driven by the dyads’ synchronous breathing^54^ and indicates distinct mechanisms driving cardiac and respiratory synchrony on a group level: dyadic respiratory synchrony was closely linked to auditory–motor coordination, while dyadic cardiac synchrony may have arisen independently of overt behavioral synchrony.

Finally, individual differences observed in the randomly paired dyad members shaped the dyad’s auditory-motor and cardiac synchrony in the Production tasks. Previous studies have demonstrated that partners with similar Spontaneous Production Rates exhibited increased auditory-motor synchronization^48,58^. The current study replicated this finding and extended it to the physiological level: Dyadic members with similar baseline heart rates exhibited greater cardiac synchronization during joint production. An important step in understanding how intrinsic frequencies influence physiology, a dynamical systems perspective suggests that two hearts may function as coupled oscillators whose similar intrinsic frequencies require less external energy to adapt and maintain synchrony, as frequency-matched systems naturally fall into stable phase relationships with less external forcing^43,59,60^. Thus, individuals with closely matched intrinsic frequencies behaviorally or physiologically likely synchronize more efficiently due to lower energy costs in adapting to each other’s biological rhythms.

Dyadic analysis of behavioral and physiological synchrony requires different design considerations to permit comparable interpersonal synchronization measures. First, a data collection interface that enables the capture of low-latency, high-precision auditory sequences is necessary to ensure millisecond-level accuracy in measuring multiple participants’ temporal coordination^61^. Second, a perturbation paradigm must be implemented to ensure consistency across dyad members; both participants perceived or produced identical rhythmic sequences in all tasks, controlling for the possible role of perception in facilitating physiological synchrony. Third, the same perturbation cues must be presented to both participants to allow for direct comparison across perturbation conditions. Finally, causal analyses within each dependent variable (behavioral asynchronies, respiratory and cardiac measures) minimize the need for data interpolation when comparing time series based on different temporal resolution. These dyadic adjustments permit us to make robust causal inference between different types of distinct physiological and behavioral signals. Future studies may extend these design issues to larger groups of interpersonal coordination^60^.

In sum, by integrating behavioral and physiological measures within a controlled experimental framework, this study provides novel insights into how auditory-motor, respiratory, and cardiac synchrony interact and dynamically emerge within and between individuals. These findings offer a comprehensive perspective on physiological synchronization in dyads, and hold promise for future measurement of interpersonal coordination in real-world settings. For instance, strategically modulating breathing patterns or pairing individuals with compatible physiological rhythms could optimize synchrony in collaborative tasks such as musical ensembles, therapeutic interactions, or team-based activities requiring high coordination. Additionally, these insights may inform interventions for individuals with social communication challenges by maximizing physiological entrainment strategies to improve social attunement and group cohesion.

## 4. Methods

### 4.1 Participants

A total of 62 participants (14 males, 48 females, and 2 nonbinary; age range: 18–34 years; M = 22.55, SD = 4.20) were recruited in the Montreal community and randomly assigned to 31 pairs to perform the dyadic tasks. The sample size of 31 dyads was determined a priori using G*Power, with an estimated medium effect size of *f* = 0.25 and power of 0.85 to detect significant differences across the three conditions. All participants had at least six years of individual music instruction and had never performed music with each other before the study (M = 11.87 years, SD = 4.06) and provided informed consent before beginning the experiment. The study was approved by McGill University’s Research Ethics Board. Participants received either course credit or a small honorarium upon completing the experiment which lasted approximately 90 minutes

### 4.2 Materials

The auditory sequences used in the Perception and Production tasks featured repeating 16-note binary meter pattern, each containing a total of 16 beats (one beat = 500 ms): presented with a classic piano timbre preset on a Roland Studio Canvas SD-50 tone generator. One pattern contained five ascending tones (G4, A4, B4, C5, D5), three beats of silence, five descending tones (D5, C5, B4, A4, G4), and three more beats of silence. A second auditory pattern, presented one octave higher than the first pattern, was designed to be discriminable when both patterns were produced simultaneously in the Production conditions. Trials in the Production condition began with eight metronome beats at 500 ms intervals to establish the indicated rate, presented at a high-hat drum timbre on the SD-50 tone generator. Perturbation cues occurred eight times in each Production trial and were placed pseudo-randomly within 500-1000 ms after the first silent beat in each auditory pattern.

Participants completed two questionnaires: A customized musical background questionnaire and Goldsmith’s Musical Sophistication Questionnaire^62^ at the start of the experiment to determine their past experience with music and level of expertise (results see SI.12). Additionally, participants completed a Social Interaction Questionnaire^63^ after each Production condition to assess their subjective judgments about their partner interaction during the experiment on a 7-point Likert scale (results see SI. Part II).

### 4.3 Equipment

To ensure precise sequence production timing, Arduino-based force-sensitive tapping pads were used connected to a Toshiba Linux computer via MIDI^64^. Each tap on the force sensor triggered the next tone in the stimulus melody, generated by a Roland Studio Canvas SD-50 and delivered to the participant through Sennheiser HD650 headphones. Timing values were recorded with 1-millisecond resolution using FTAP^61^.

Physiological data were recorded using Biopoint sensors (Quebec, Canada) and Vernier GDX-RB respiration belts (Oregon, USA). The Biopoint sensors, placed on the non-dominant wrist, captured bipolar ECG data at 500 Hz, with sensor leads positioned on the underside of the wrist and a ground electrode attached to the dominant-side waist. The respiration belts, secured around the chest near the diaphragm, recorded inhalation and exhalation motion at 10 Hz. All data streams were synchronized using custom Python scripts on a Linux computer.

### 4.4 Design and Procedure

The within-subject design consisted of four separate tasks performed in the following order: Individual differences measurement, Baseline measurement, Perception task, and Production task (Figure 1B).

After participants provided consent, they completed the two music background questionnaires. Then the participants completed a standardized individual Spontaneous Rate (SPR) measurement task individually, to determine the rate of their spontaneous productions of a familiar melody^35,36,41,48^. Participants first confirmed their familiarity with the melody “Twinkle Twinkle Little Star”, an isochronous rhythm used in previous studies to measure participants’ natural frequencies^35,41,48^. Participants produced the melody by tapping on the force-sensitive pad with their index finger at a steady and consistent rate, where each tap produced the next tone in the sequence, heard over headphones in a marimba timbre (GM2, patch 13, bank #0) produced on a SD-50 tone generator at a fixed loudness level. They were instructed to tap the melody for three iterations without any pauses on each trial and they completed 3 trials. Each participant’s spontaneous rate was calculated from the middle two complete repetitions of the melody to capture maximally stable rhythmic behavior (excluding the initial 16 and final 32 tones)^35,36^. The mean and standard error of the intertap intervals (ITI) of the produced melodies on each trial were computed as each participants’ Spontaneous rate.

Next, the participants were brought to the same testing room to complete the rest of the tasks. First, a ninety-second-long silent Baseline measurement was conducted to assess participants’ resting heart rate and respiratory patterns during which, participants were asked to sit quietly and remain still.

The Perception task was conducted to assess physiological responses during melody perception and to ensure that participants could reliably detect both melodies when presented simultaneously and the perturbation cues. During the Normal Perception trial, participants listened to the melody without perturbation cues; during the Perturbation Perception trial, perturbation cues were embedded at the same locations as in the Production task. Participants were told that some trials would contain high-pitched cues and they were instructed to count the number of cues on each trial. There was a total of three trials. Participants were required to report the correct number of cues to continue in the study.

Next, the Production task was conducted; participants were instructed to produce the melody on the force sensor pads while they were seated face-to-face, with a visual barrier preventing them from seeing each other’s hands (Figure 1A). They were asked to synchronize their melody with their partner as accurately as possible after the initial metronome cue ended, while maintaining the metronome’s rate. The Production task consisted of three conditions: Normal, Auditory, and Respiratory. The Normal condition was conducted twice (at the beginning and end of the Production task), and the order of the Auditory and Respiratory conditions was counterbalanced across pairs. Each condition included practice trials for participants to familiarize themselves with the task.

During the Normal trials, participants were instructed to produce the melody in synchrony with their partner. During the Respiratory trials, participants were instructed to perform the same melody, and that during unpredictable silent beats, a high-pitched sound would signal both participants to take a quick, deep breath. During the Auditory condition, participants were instructed to perform the same melody, and that during unpredictable silent beats, a high-pitched sound would signal both participants to immediately restart the melody on the next beat, starting with the first melody tone.

### 4.5 Data Analyses

#### 4.5.1 Behavioral data analysis

The melody production data was processed using a Python script that extracted tone onsets from each force sensor. Each participant’s inter-tap intervals (ITIs), defined as the time intervals between consecutive taps (in ms), and the dyad’s absolute asynchronies (defined as the absolute value of Participant 1’s tone onsets - Participant 2’s tone onsets) for tones intended as simultaneous (ie, from the same serial position in the melodies) were computed. Trials containing three or more ITIs exceeding 1000 ms (i.e., longer than two reference beats) or asynchronies exceeding 1000 ms were excluded from the analysis. In total, 14 trials (3.8% of all trials) and 43 asynchronies (0.1% of all asynchrony data) were excluded.

#### 4.5.2 Physiological data analysis

Both respiratory and cardiac data were preprocessed and cleaned using NeuroKit2 in Python 3.8 following established procedures^65^. Cardiac data quality was first assessed using an automated algorithm^66^, and only the signals classified as high-quality were included in the analyses. No trials were excluded based on this assessment. The raw ECG signals were then preprocessed using a fifth-order, 0.5 Hz high-pass Butterworth filter, followed by powerline filtering^65^. R-R intervals were extracted using the NeuroKit algorithm and corrected for potential artifacts^67^. To ensure data length consistency between participants^7,18^, R-R intervals were resampled to 0.2-second intervals BPM data^68^.

Respiratory data were normalized, detrended, and filtered using a second-order Butterworth bandpass filter (0.05–3 Hz)^69,70^. Then the respiration peak detection was performed by the NeuroKit2 software; Inhale-to-inhale intervals were then extracted from the cleaned respiration signals^69^. To confirm compliance with the deep breathing instructions in the Respiratory condition, we visually inspected the alignment between breathing cycles and perturbation cue locations. Trials were excluded if fewer than six of the eight expected deep breaths were detected as prominent respiration cycles. Based on this criterion, 8 trials (2.1% of the total trials) were removed from the dataset.

Physiological synchronization was assessed using Phase Locking Value (PLV) in a Python 3.8 script; PLV is a measure that quantifies the consistency of phase differences between two time series^49,50^, and has been widely applied to two periodic time series to quantify the level of synchronization in their relative phase^12,47,49,50,71,72^. PLV provides a normalized value ranging from 0 (no synchronization) to 1 (perfect synchronization), indicating the stability of the phase relationship over time and, consequently, the strength of synchronization.

We applied the Hilbert transform to extract the instantaneous phase of each participant’s time series at each time point. For dyadic cardiac and respiratory PLV analyses, the phase difference between the two pre-processed time series (Participant 1’s heart rate/respiration – Participant 2’s heart rate/respiration) was computed at each time point as Δ□(*k*). For within-individual cardiorespiratory PLV analyses, to account for frequency and sampling rate differences between cardiac and respiratory signals, the R peaks timing was extracted from ECG signals, and mapped with respiratory signals, n:m (n = cardiac frequency, m = respiratory frequency) phase difference was calculated as: Δ*ϕ*(*k*) = *n* · *ϕ_resp_*(*k*) - *m* · *ϕ_ecg_*(*k*). Following established procedures^45,47^, m was fixed at 1, and n was determined by rounding the ratio of cardiac to respiratory frequency to the nearest integer for every trial. PLV for each trial was calculated using the following equation:

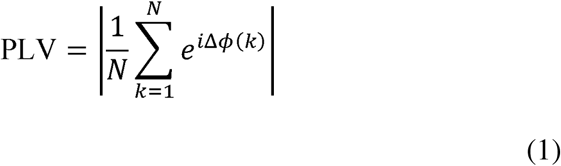

where N is the number of time points in the trial. The magnitude of the mean resultant vector reflects the consistency of phase differences within one trial, with higher PLV values indicating stronger phase-locking between signals.

#### 4.5.3 Statistical analysis

All statistical analyses were conducted in R (RStudio 2024.10) using ANOVA with repeated measures for comparisons across Baseline, Perception, and Production conditions. A second set of ANOVAs compared differences across the 3 Production conditions (Normal, Auditory, Respiratory). When a significant main effect was observed (α = 0.05), Holm–Bonferroni– corrected post hoc tests were applied to adjust for multiple pairwise comparisons. To examine relationships within and between behavioral and physiological measures, Pearson correlation analyses were performed; Spearman rho was performed when data did not meet normality assumptions. Outliers—defined as data points exceeding three standard deviations from the mean—were excluded from the correlation analysis (only one data point got removed). PLV values were computed separately for each trial in the Normal, Auditory, and Respiratory conditions to assess how experimental manipulations influenced cardiac and respiratory synchronization. A repeated-measures analysis of variance (ANOVA) was conducted to compare PLV values across conditions, testing whether task demands and perturbation cues modulated physiological coupling between participants.

## Acknowledgments

The authors acknowledge the assistance of Elizabeth Harrigan, Kai Mikkelsen, and Joshua Samuels in data collection and technical assistance. This work was partially funded by grants from the Natural Sciences and Engineering Research Council of Canada—Create Graduate Fellowship to W.Y., and from the Natural Sciences and Engineering Research Council of Canada (Discovery Grant 298173) and Canada Research Chair to C.P.

