## Supplementary Information for "Respiratory synchrony and individual differences causally influence dyadic interpersonal coordination"

PREPRINT – Do not quote

May 2025

**1.** To determine whether the perturbation effects on the tone onset asynchronies only affected tone synchronization immediately after the perturbation cue in the Production conditions, we removed the first (SI Table 1) plus the second (SI Table 2) asynchronies following each cued location from the data and repeated the analysis. The repeated measures ANOVA results showed the same pattern of results, with larger asynchronies in Respiratory, then Auditory, then Normal performance conditions.

**SI Table 1. Tone onset asynchronies excluding one location following each cue**

| *Within Subjects Effects* | | | | | | | | | | | | | | | | |
| --- | --- | --- | --- | --- | --- | --- | --- | --- | --- | --- | --- | --- | --- | --- | --- | --- |
| Cases | | | Sum of Squares | | df | Mean Square | | | | | F | p | | η² | | |
| Production tasks |  | | 2360.099 |  | 2 |  | 1180.049 | |  | | 8.870 |  | < .001 | |  | 0.228 |
| Residuals |  | | 7982.362 |  | 60 |  | 133.039 | |  | |  |  |  | |  |  |
| *Note.*  Type III Sum of Squares | | | | | | | | | | | | | | | | |
| *Post Hoc Comparisons - Production tasks* | | | | | | | | | | | | | | | | |
|  | |  | | Mean Difference | | | | SE | | | t | Cohen's d | | | p_holm_ | |
| Normal |  | Auditory |  | -4.043 | | |  | 1.883 | |  | -2.147 |  | -0.228 | |  | 0.045 |
|  |  | Respiratory |  | -12.118 | | |  | 3.308 | |  | -3.663 |  | -0.684 | |  | 0.003 |
| Auditory |  | Respiratory |  | -8.075 | | |  | 3.356 | |  | -2.406 |  | -0.456 | |  | 0.045 |
| *Note.*  P-values adjusted for Holm's comparisons among 3 means | | | | | | | | | | | | | | | | |

**SI Table 2. Tone onset asynchronies excluding two locations following each cue**

| *Within Subjects Effects* | | | | | | | | | | | | |
| --- | --- | --- | --- | --- | --- | --- | --- | --- | --- | --- | --- | --- |
| Cases | | Sum of Squares | | df | | Mean Square | | F | | p | | η² |
| Production task |  | 1743.757 |  | 2 |  | 871.879 |  | 7.521 |  | 0.001 |  | 0.200 |
| Residuals |  | 6955.596 |  | 60 |  | 115.927 |  |  |  |  |  |  |
| Note.  Type III Sum of Squares | | | | | | | | | | | | |

| *Post Hoc Comparisons - Condition* | | | | | | | | | | | | |
| --- | --- | --- | --- | --- | --- | --- | --- | --- | --- | --- | --- | --- |
|  | |  | | Mean Difference | | SE | | t | | Cohen's d | | p_holm_ |
| Normal |  | Auditory |  | -3.977 |  | 1.915 |  | -2.077 |  | -0.236 |  | 0.088 |
|  |  | Respiratory |  | -10.504 |  | 3.022 |  | -3.476 |  | -0.630 |  | 0.005 |
| Auditory |  | Respiratory |  | -6.527 |  | 3.105 |  | 201 |  | -0.394 |  | 0.088 |
| *Note.*  P-values adjusted for Holm's comparisons among 3 means | | | | | | | | | | | | |

**2.** A repeated measures ANOVA on the mean inter-tap intervals (ITIs) produced in the Production conditions showed that participants produced the melodies significantly slower in the Respiratory condition than in the Normal and Auditory conditions, as reflected in the largest mean ITI (SI Table 3). This effect persisted after removing the first ITI value following each cue (SI Table 4), suggesting that the respiratory perturbations slowed down the dyads’ production rates in the synchronization task.

**SI Table 3. Mean inter-tap intervals of conditions:**

| *Within Subjects Effects* | | | | | | | | | | |
| --- | --- | --- | --- | --- | --- | --- | --- | --- | --- | --- |
| Cases | | Sum of Squares | | df | | Mean Square | | F | | p |
| Condition |  | 2350.913 |  | 2 |  | 1175.457 |  | 10.665 |  | < .001 |
| Residuals |  | 6621.837 |  | 60 |  | 110.215 |  |  |  |  |
| Note.  Type III Sum of Squares | | | | | | | | | | |

| *Post Hoc Comparisons - Condition* | | | | | | | | | | | | |
| --- | --- | --- | --- | --- | --- | --- | --- | --- | --- | --- | --- | --- |
|  | |  | | Mean Difference | | SE | | t | | Cohen's d | | p_holm_ |
| Normal |  | Auditory |  | 0.514 |  | 2.335 |  | 0.220 |  | 0.019 |  | 0.827 |
|  |  | Respiratory |  | -10.399 |  | 2.447 |  | -4.250 |  | -0.390 |  | < .001 |
| Auditory |  | Respiratory |  | -10.913 |  | 3.145 |  | -3.470 |  | -0.409 |  | 0.003 |
| Note.  P-value adjusted for comparing a family of 3 | | | | | | | | | | | | |

**SI Table 4. Mean inter-tap intervals of conditions (removed the first ITI value after each cue):**

| *Within Subjects Effects* | | | | | | | | | | |
| --- | --- | --- | --- | --- | --- | --- | --- | --- | --- | --- |
| Cases | | Sum of Squares | | df | | Mean Square | | F | | p |
| Condition |  | 1372.344 |  | 2 |  | 686.172 |  | 7.447 |  | 0.001 |
| Residuals |  | 5528.470 |  | 60 |  | 92.141 |  |  |  |  |
| Note.  Type III Sum of Squares | | | | | | | | | | |

| *Post Hoc Comparisons - Condition* | | | | | | | | | | | | |
| --- | --- | --- | --- | --- | --- | --- | --- | --- | --- | --- | --- | --- |
|  | |  | | Mean Difference | | SE | | t | | Cohen's d | | p_holm_ |
| Normal |  | Auditory |  | 0.534 |  | 2.210 |  | 0.241 |  | 0.020 |  | 0.811 |
|  |  | Respiratory |  | -7.869 |  | 2.132 |  | -3.692 |  | -0.299 |  | 0.003 |
| Auditory |  | Respiratory |  | -8.402 |  | 2.899 |  | -2.898 |  | -0.319 |  | 0.014 |
| Note.  P-value adjusted for comparing a family of 3 | | | | | | | | | | | | |

**3.** Paired sample t-tests were conducted to test the physiological activity differences between the first and last Normal Production conditions for each of the mean respiration phase locking values and mean cardiac phase locking values. Both indices showed no significant differences between the first and last Normal Production conditions (SI Table 5, SI Table 6).

**SI Table 5. Comparison of respiration PLV between Normal-first and Normal-last Production conditions.**

| *Paired Samples T-Test* | | | | | | | | | | |
| --- | --- | --- | --- | --- | --- | --- | --- | --- | --- | --- |
| Normal-first | |  | | Normal-last | | t | | df | | p |
| 0.665 |  | - |  | 0.739 |  | -1.506 |  | 30 |  | 0.143 |
| Note.  Student's t-test. | | | | | | | | | | |

**SI Table 6. Comparison of cardiac PLV between Normal-first and Normal-last Production conditions.**

| *Paired Samples T-Test* | | | | | | | | | | |
| --- | --- | --- | --- | --- | --- | --- | --- | --- | --- | --- |
| Normal-first | |  | | Normal-last | | t | | df | | p |
| 0.187 |  | - |  | 0.186 |  | 0.023 |  | 30 |  | 0.982 |
| Note.  Student's t-test. | | | | | | | | | | |

**4.** Paired t-tests were conducted to test physiological differences between the Perception–normal (melody without perturbation cue) and Perception–perturbation (melody without perturbation cue) conditions. Both indices, including respiration phase locking values (SI Table 7) and cardiac phase locking values (SI Table 8), indicated no significant differences.

**SI Table 7. Comparison of respiration PLV between Perception-Normal and Perception-Perturbation conditions.**

| *Paired Samples T-Test* | | | | | | | | | | |
| --- | --- | --- | --- | --- | --- | --- | --- | --- | --- | --- |
| Perception-Normal | |  | | Perception-Perturbation | | t | | df | | p |
| 0.589 |  | - |  | 0.591 |  | -0.51 |  | 30 |  | 0.959 |
| Note.  Student's t-test. | | | | | | | | | | |

**SI Table 8. Comparison of cardiac PLV between Perception-Normal and Perception-Perturbation conditions.**

| *Paired Samples T-Test* | | | | | | | | | | |
| --- | --- | --- | --- | --- | --- | --- | --- | --- | --- | --- |
| Perception-Normal | |  | | Perception-Perturbation | | t | | df | | p |
|  | 0.205 | - |  |  | 0.164 | 1.408 |  | 30 |  | 0.170 |
| Note.  Student's t-test. | | | | | | | | | | |

**5. Surrogate Analyses for dyadic cardiac and respiratory synchrony**

To evaluate whether cardiac and respiratory synchrony arise specifically from dyadic interactions and are modulated by experimental conditions, we conducted two surrogate analyses across Production tasks. In the first analysis, we tested whether the observed synchrony reflected genuine interpersonal coupling, rather than shared responses to common external stimuli. To this end, Participant 1 from each dyad was reassigned to a randomly selected, non-original partner from a different dyad within the same condition (e.g., Participant 1 from Pair 1 in the Normal condition was reassigned to Participant 2 from each other dyad in the Normal condition). Surrogate phase-locking values (PLVs) were then computed for both cardiac and respiratory signals. We implemented a Monte Carlo simulation with 1,000 iterations, in which surrogate PLVs were randomly sampled while preserving the original dyad-condition structure. This approach enabled us to generate a null distribution of mean differences expected in the absence of true physiological synchrony. The empirical p-value was defined as the proportion of iterations in which the mean PLV from real dyads was less than or equal to that from surrogate dyads (i.e. Null hypothesis: real PLV – surrogate PLV = 0). Results rejected the null hypothesis and revealed that real dyadic respiratory PLVs (N = 352) were significantly higher than those derived from surrogate pairs (N = 59936, observed mean difference = 0.05, *p* = 0.0042, SI. Figure 1). Similarly, real dyadic cardiac PLVs (N = 352) were significantly greater than surrogate values (N = 59936, observed mean difference = 0.03, *p* = 0.019, SI. Figure 2).

Second, we conducted a within-dyad surrogate analysis across the three Production conditions (Normal, Auditory, and Respiratory) to assess whether physiological PLVs varied as a function of experimental manipulation. For each dyad, we paired the time series of Participant 1 from one condition (e.g., Normal) with the time series of Participant 2 from a different condition within the same dyad to generate surrogate PLVs. This approach enabled us to test whether the observed synchrony was specifically driven by the experimental condition, rather than reflecting stable interpersonal dynamics or confounding artifacts unrelated to task context. A similar Monte Carlo simulation with 1,000 iterations revealed that respiratory PLVs from the original condition pairings (N = 352) were significantly higher than those from reassigned conditions (N = 4612, mean difference = 0.07, p < 0.001, SI. Figure 3). Cardiac PLVs (N = 352) from the original data were also significantly greater than their surrogate counterparts (N = 4612, mean difference = 0.04, p < 0.001, SI. Figure 4), supporting the interpretation that the observed physiological synchrony was condition-dependent.

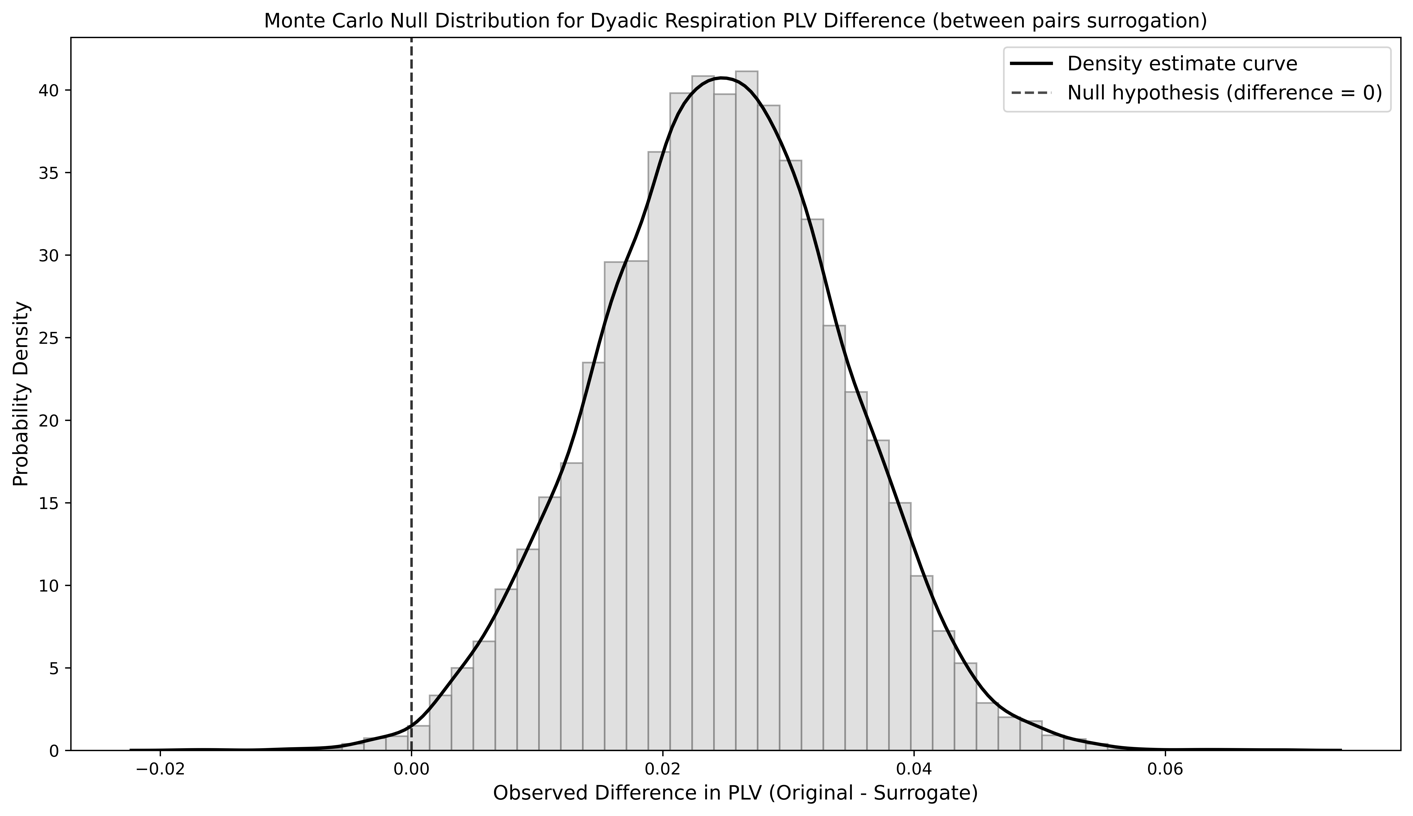

**SI. Figure 1.** Null distribution for dyadic respiratory PLV difference between real dyads PLV – surrogate dyads PLV, dashed line indicate the null hypothesis (PLV difference = 0).

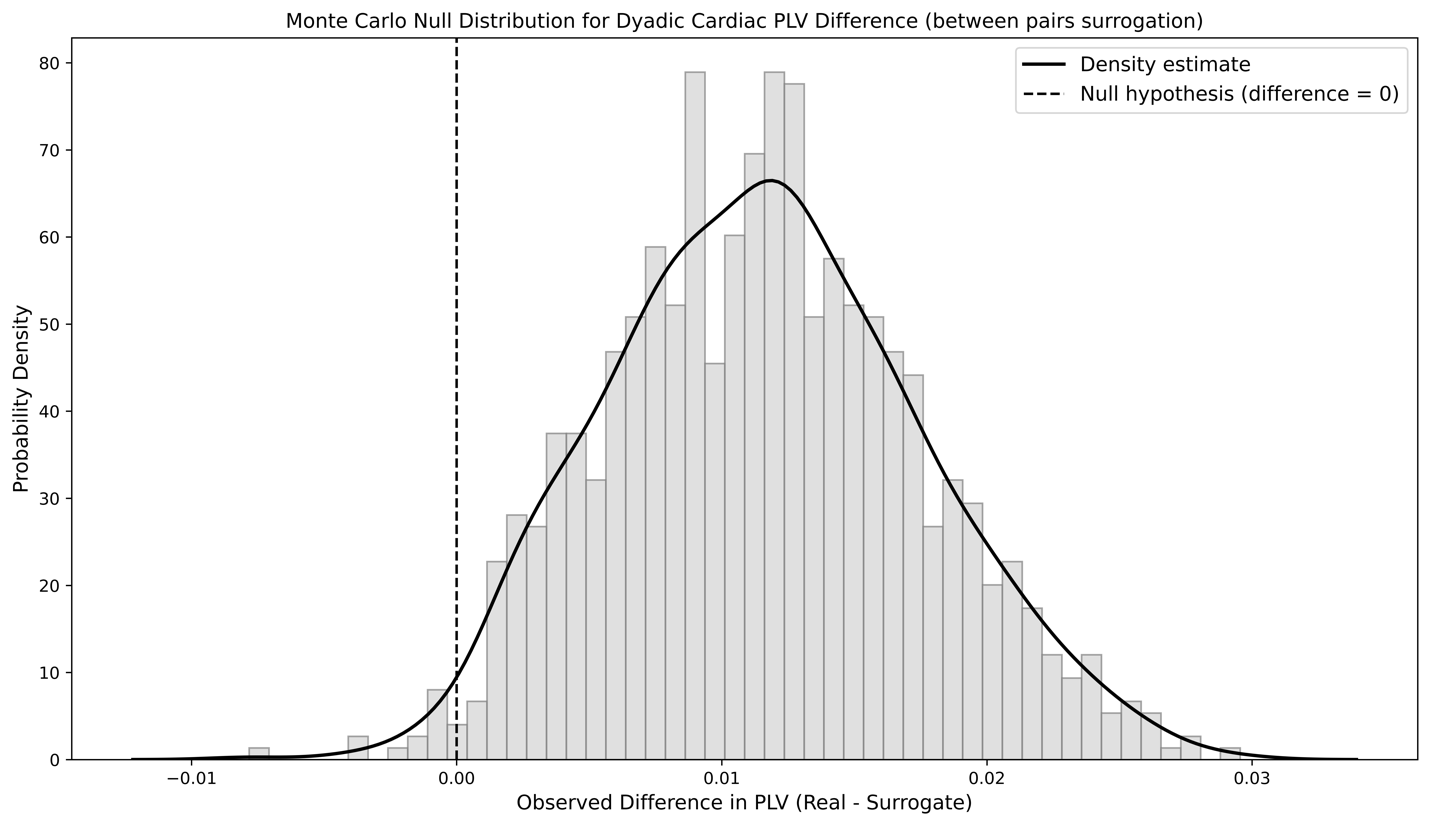

**SI. Figure 2.** Null distribution for dyadic cardiac PLV difference between real dyads PLV – surrogate dyads PLV, dashed line indicate the null hypothesis (PLV difference = 0).

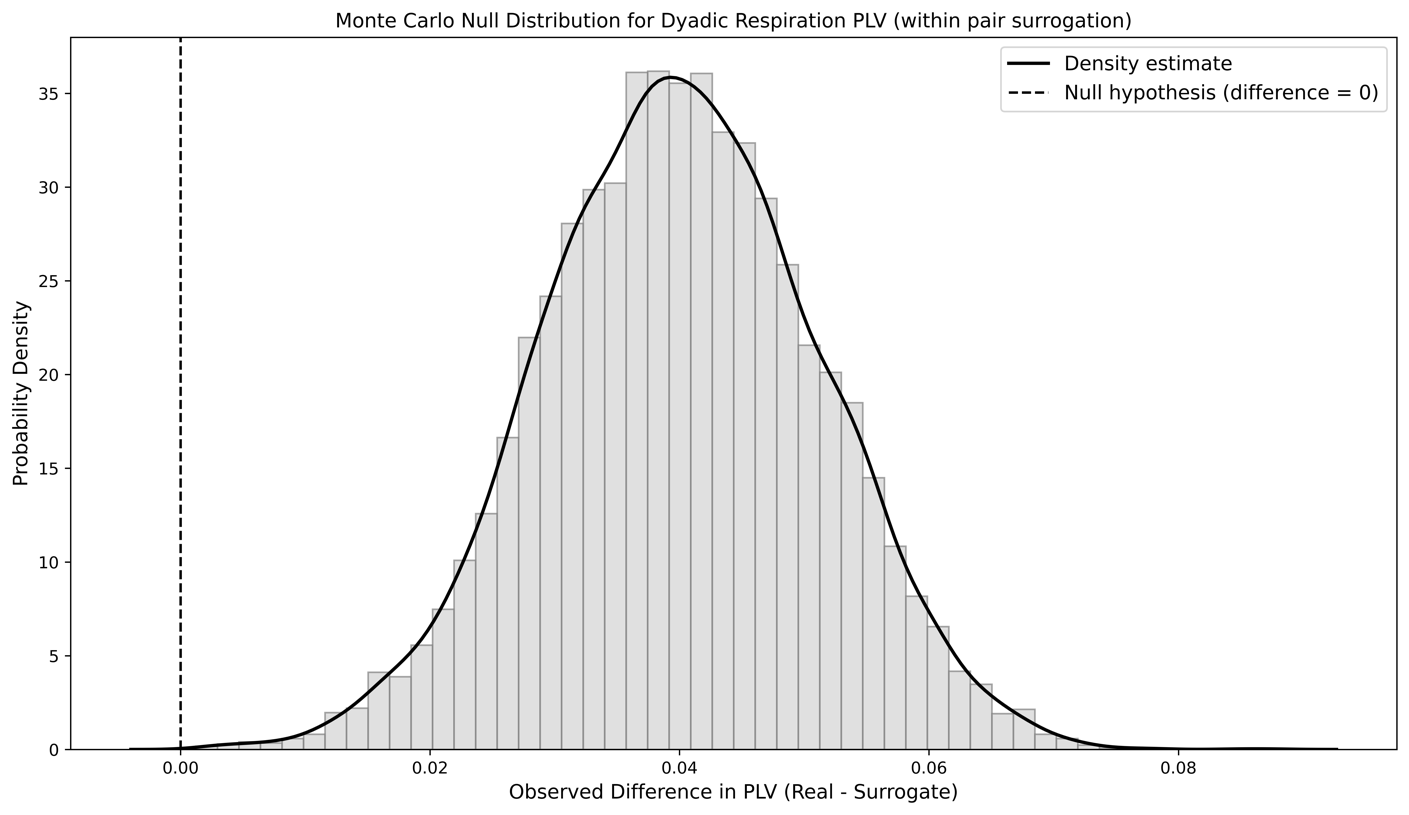

**SI. Figure 3.** Null distribution for dyadic respiratory PLV difference between original condition generated PLV – reassigned condition generated dyads PLV, dashed line indicate the null hypothesis (PLV difference = 0).

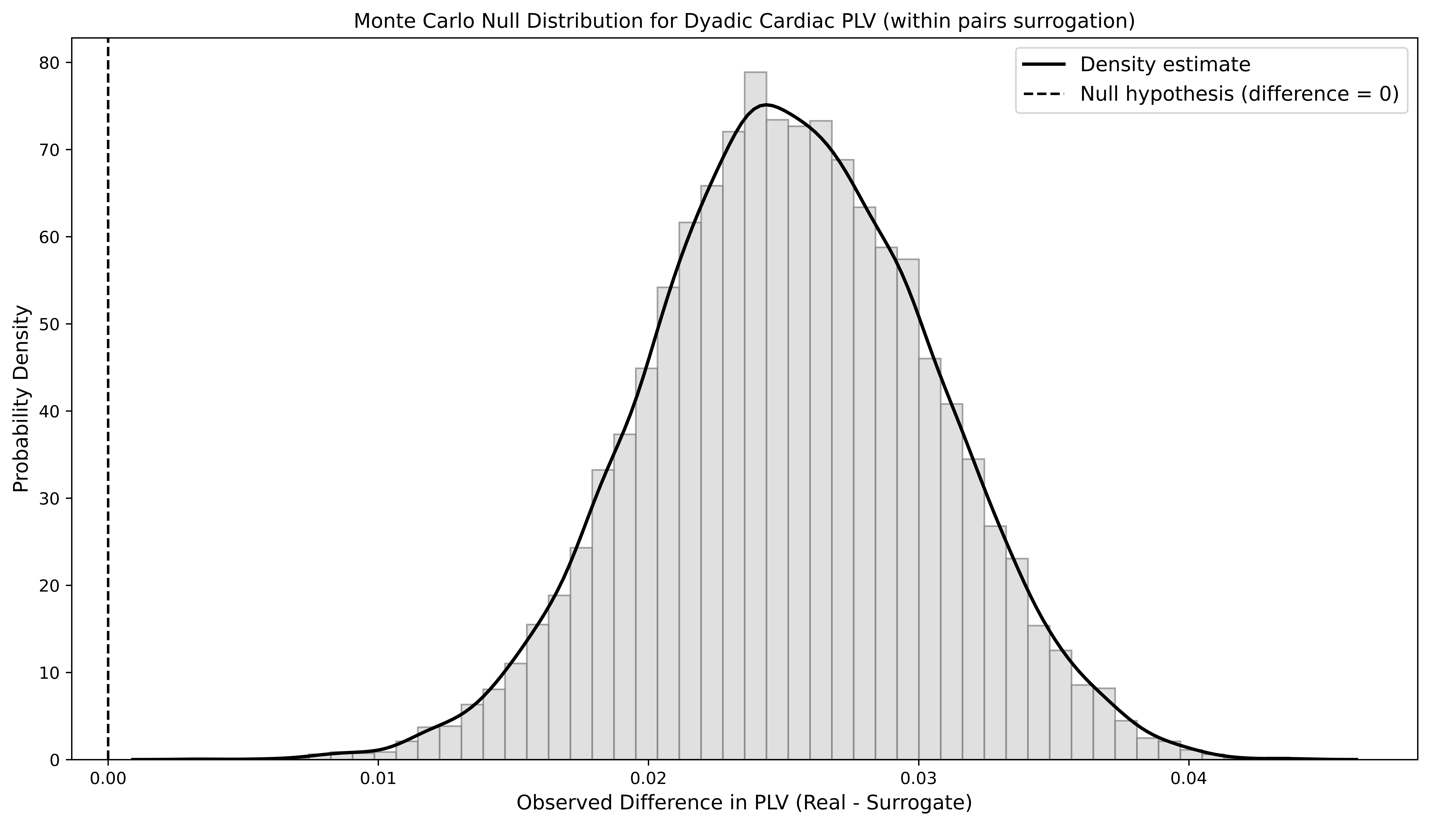
 **SI. Figure 4.** Null distribution for dyadic cardiac PLV difference between original condition generated PLV – reassigned condition generated dyads PLV, dashed line indicate the null hypothesis (PLV difference = 0).

**6.** To further test the correlation between the dyad’s mean resting heart rate differences and their mean cardiac PLV during Baseline and Perception tasks, the Spearman correlation has been conducted between the mean resting heart rate differences and mean PLV. The correlations were not significant in the Baseline or Perception tasks (SI Table 9).

**SI Table 9. Correlation between dyads' resting heart rate differences (Baseline) and cardiac PLV during Baseline and Perception tasks.**

| Variables | | Spearman's rho | | p-value | |
| --- | --- | --- | --- | --- | --- |
| Baseline HR Diff - Baseline Cardiac PLV |  |  | 0.097 |  | .602 |
| Baseline HR Diff – Perception Cardiac PLV |  |  | -0.185 |  | .317 |

In addition, no significant correlation was observed between differences in partners' resting respiration rates (measured in mean inhale-inhale interval differences) and their respiratory synchronization (Spearman *r* = 0.290, *p* = 0.114). The lack of significant associations between these individual differences and respiratory synchrony suggests that distinct mechanisms govern cardiac and respiratory synchronization in dyadic interactions.

**7.** To test the correlation between the dyads' cardiac and auditory-motor synchronization, we conducted Pearson correlations between mean cardiac phase locking values and mean onset absolute asynchronies in Normal, Auditory, and Respiratory conditions. There was no significant correlation between cardiac PLVs and tone-onset asynchronies (SI Table 10).

**SI Table 10. Correlation between mean cardiac PLV and mean absolute asynchronies in Normal, Auditory, and Respiratory conditions**

| *Pearson Correlations - Normal condition* | | | | |
| --- | --- | --- | --- | --- |
| Variables |  | | |  |
| Mean absolute asynchronies – Mean cardiac PLV |  | Pearson r |  | -0.096 |
|  |  | p-value |  | 0.609 |
| *Pearson Correlations* - Respiratory condition | | | | |
| Variables |  | | |  |
| Mean absolute asynchronies – Mean cardiac PLV |  | Pearson r |  | 0.051 |
|  |  | p-value |  | 0.784 |

| *Pearson Correlations* - Auditory condition | | | | | |
| --- | --- | --- | --- | --- | --- |
| Variables |  | |  | | |
| Mean absolute asynchronies – Mean cardiac PLV |  | Pearson r | |  | 0.161 |
|  |  | p-value | |  | 0.386 |

**8.** To test the relationship between the dyads' cardiac and respiration synchronization, we conducted Pearson correlations between mean cardiac PLVs and mean respiration PLVs in Normal, Auditory, and Respiratory conditions across dyads. No significant correlations were observed in any of the three conditions (SI Table 11).

**SI Table 11. Correlation between mean cardiac PLVs and mean respiration PLVs in Production conditions**

| *Pearson Correlations* - Normal condition | | | | |
| --- | --- | --- | --- | --- |
| Variables | |  | |  |
| Mean cardiac PLVs – Mean resp PLVs |  | Pearson r |  | -0.098 |
|  |  | p-value |  | 0.602 |

| *Pearson Correlations* - Auditory condition | | | | |
| --- | --- | --- | --- | --- |
| Variables | |  | |  |
| Mean cardiac PLVs – Mean resp PLVs |  | Pearson r |  | 0.047 |
|  |  | p-value |  | 0.804 |

| *Pearson Correlations* - Respiratory condition | | | | |
| --- | --- | --- | --- | --- |
| Variables | |  | |  |
| Mean cardiac PLVs – Mean resp PLVs |  | Pearson r |  | 0.260 |
|  |  | p-value |  | 0.159 |

**9. Surrogate Analyses for individual cardiorespiratory synchrony**

To assess whether the n:m phase-locking value reflects genuine within-individual cardiorespiratory phase coupling, we conducted a surrogate analysis in which each individual’s cardiac signal was paired with the respiration signal of every other participant in the same Production conditions. For each surrogate pair, the corresponding cardiac-to-respiratory frequency ratio was calculated to determine the appropriate n:m ratio for each trial, respectively, which was then applied to compute surrogate PLVs. This procedure generated a distribution of surrogate cardiorespiratory PLVs for comparison against the original (self) pairings. We then implemented a Monte Carlo simulation with 1,000 iterations, in which surrogate PLVs were randomly sampled while preserving the original individual–condition structure. The empirical p-value was defined as the proportion of iterations in which the mean PLV from the genuine cardiorespiratory pairings was less than or equal to that from the surrogate combinations (i.e. Null hypothesis: real PLV – surrogate PLV = 0). The results showed that real individual-level cardiorespiratory PLVs were significantly higher than those derived from surrogate pairings (mean difference = 0.024, *p* < 0.001, SI. Figure 5), supporting the specificity of within-individual phase coupling between cardiac and respiratory signals.

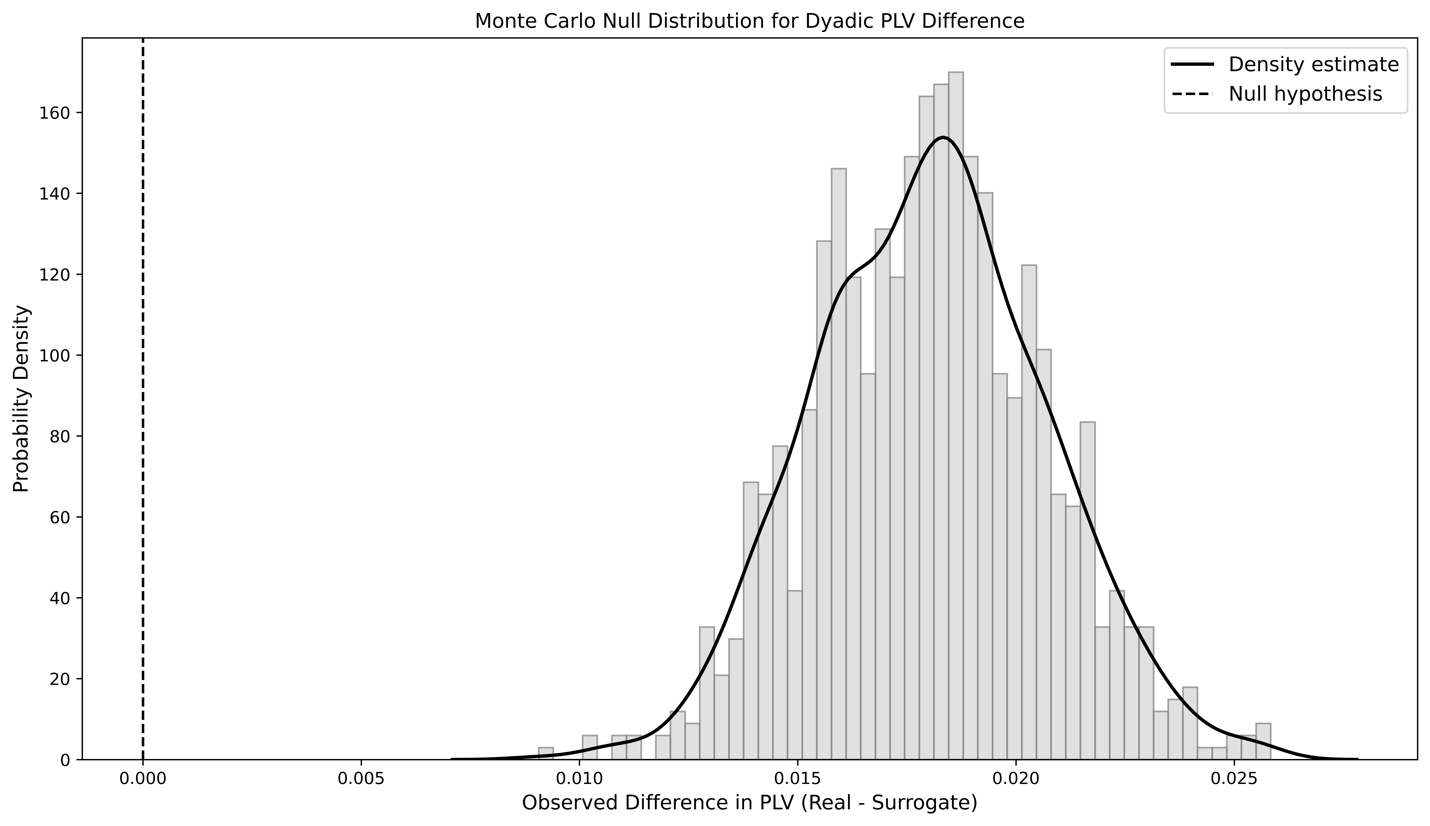

**SI. Figure 5.** Null distribution for individual cardiorespiratory PLV difference between real PLV – reassigned PLV, dashed line indicate the null hypothesis (PLV difference = 0).

**10.** Each participant's mean heart rate (in BPM) was first compared across the Baseline, Perception, and Production tasks. A repeated-measures ANOVA showed no significant differences *F*(2, 60) = 1.18, *p* = .316. Since there are no significant differences between the first Normal and last Normal conditions *t*(30)=-1.426, *p* = 0.164, the two normal conditions have been averaged to one condition. Mean heart rate across the Normal, Auditory, and Respiratory perturbation conditions differed significantly, *F*(2, 60) = 6.439, *p* = .003, *η²* = .177 (SI Figure 6). Post-hoc comparisons revealed that the mean heart rate was significantly higher in the Respiratory condition than in the Normal condition, *t*(30) = -3.046, *p_holm_* = .014, *d* = -0.337, and the Auditory condition, *t*(30) = -2.455, *p_holm_* = .004, *d* = -0.239. There was no significant difference in mean heart rate between the Normal and Auditory conditions, *t*(30) = -1.238, *p_holm_* = 0.225.

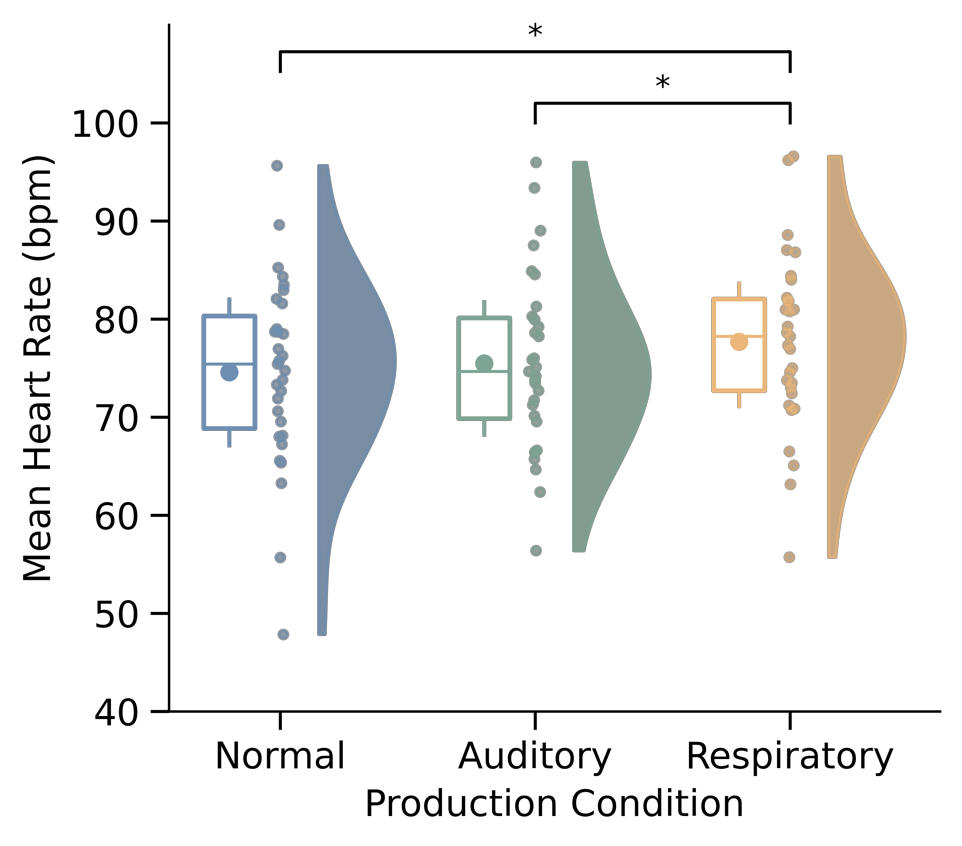

**SI. Figure 6. Individual Mean Heart Rates (in BPM) in three Production conditions:** Box plots display medians (horizontal lines) and means (bold points). Significance markers: p < 0.05 (*), p < 0.01 (****),** p **< 0.001 (*****).

**11.** Each participant's mean respiration rate (in BPM) was first compared across the Baseline, Perception, and Production tasks. There were no significant differences among the three tasks, *F*(2,60) = 1.037, *p* = 0.361. Since there is a significant difference between the mean respiration rate between Normal-first and Normal-last conditions (*t*(30) = 3.171, *p* = 0.003), we test the difference in mean respiration rates between Normal-first, Auditory, Respiratory, and Normal-last conditions. However, a significant difference was found among the first Normal, Auditory, Respiratory, and second Normal conditions, *F*(3,90) = 108.692, *p* < .001, *η²* = .78 (SI Figure 7). Post hoc tests revealed that the respiration rate in the Respiratory condition was significantly slower than in all other conditions, confirming the effect of respiratory perturbations as deep breathing slows respiration. The remaining conditions (Normal and Auditory) showed no significant differences in individual respiration rates, the post hoc test results are shown below (SI Table 12):

**SI Table 12. Mean respiration rates in Normal-first, Auditory, Respiratory, and Normal-last conditions.**

| *Post Hoc Comparisons - Conditions* | | | | | | | | | | | | | | |
| --- | --- | --- | --- | --- | --- | --- | --- | --- | --- | --- | --- | --- | --- | --- |
|  | |  | | Mean Difference | | SE | | df | | t | | Cohen's d | | p_holm_ |
| Normal-first |  | Auditory |  | 0.238 |  | 0.223 |  | 30 |  | 1.066 |  | 0.098 |  | 0.295 |
|  |  | Respiratory |  | 7.014 |  | 0.608 |  | 30 |  | 11.545 |  | 2.898 |  | < .001 |
|  |  | Normal-last |  | 0.780 |  | 0.246 |  | 30 |  | 3.171 |  | 0.322 |  | 0.010 |
| Auditory |  | Respiratory |  | 6.776 |  | 0.636 |  | 30 |  | 10.652 |  | 2.800 |  | < .001 |
|  |  | Normal-last |  | 0.542 |  | 0.245 |  | 30 |  | 2.213 |  | 0.224 |  | 0.069 |
| Respiratory |  | Normal-last |  | -6.233 |  | 0.545 |  | 30 |  | -11.434 |  | -2.576 |  | < .001 |
| Note.  P-value adjusted for comparing a family of 6 estimates. | | | | | | | | | | | | | | |

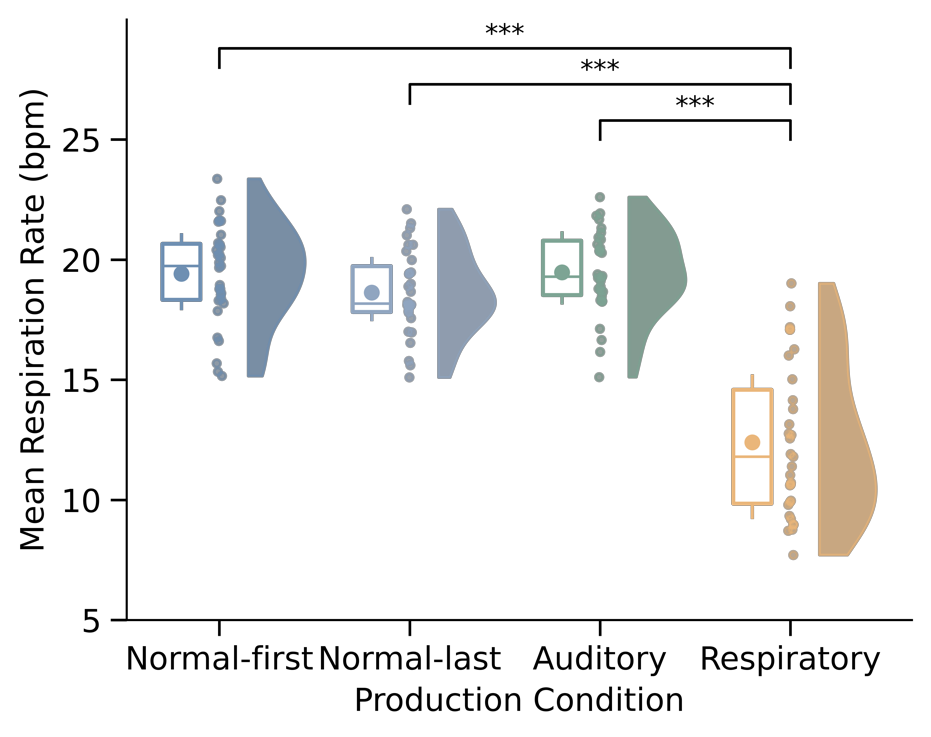

**SI. Figure 7. Individual Mean Respiration Rates (in BPM) in three Production conditions:** Box plots display medians (horizontal lines) and means (bold points). Significance markers: p < 0.05 (*), p < 0.01 (****),** p **< 0.001 (*****).

**12. Participants' musical background**

Participants' musicianship was assessed with the Goldsmith Musicianship Questionnaire, and their years of private instrument training received on their longest-duration instrument was measured.

Significant negative correlations were found between the dyads' expertise levels (measured by mean General Musical Sophistication score) and their cardiac synchronization levels (measured by mean cardiac phase locking values) in the Normal Production condition (*r* = -0.408, *p* = 0.023,SI Table 13); no significant correlation was found in Auditory and Respiratory conditions (SI Table 13). Moreover, there were no significant correlations between dyads' mean respiratory PLVs and the dyads' mean General Musical Sophistication scores in Normal, Auditory, and Respiratory conditions (SI Table 14). Moreover, no significant correlations were found between dyads' mean auditory-motor absolute asynchronies and the dyads' mean General Musical Sophistication scores in Normal, Auditory, and Respiratory conditions (SI Table 15).

Additionally, no significant correlation was found between participants’ years of private instrument training on the longest instrument and their cardiac (SI Table 16), respiratory (SI Table 17), and auditory-motor synchronization (SI Table 18) levels in Normal, Auditory, and Respiratory conditions.

| **SI. Table 13 - Correlation between mean cardiac PLVs and mean General Musical Sophistication scores** | | | | | |
| --- | --- | --- | --- | --- | --- |
| Variables |  | | |  | |
| Mean cardiac PLVs (Normal condition) – Mean General Musical Sophistication scores |  | Pearson's r |  | | -0.408 |
|  |  | p-value |  | | 0.023 |
| Mean cardiac PLVs (Auditory condition) – Mean General Musical Sophistication scores |  | Pearson's r |  | | -0.226 |
|  |  | p-value |  | | 0.222 |
| Mean cardiac PLVs (Respiratory condition) – Mean General Musical Sophistication scores |  | Pearson's r |  | | -0.099 |
|  |  | p-value |  | | 0.596 |

| **SI. Table 14 - Correlation between mean respiratory PLVs and mean General Musical Sophistication scores** | | | | | |
| --- | --- | --- | --- | --- | --- |
| Variables |  | | |  | |
| Mean respiratory PLVs (Normal condition) – Mean General Musical Sophistication scores |  | Pearson's r |  | | 0.176 |
|  |  | p-value |  | | 0.344 |
| Mean respiratory PLVs (Auditory condition) – Mean General Musical Sophistication scores |  | Pearson's r |  | | 0.136 |
|  |  | p-value |  | | 0.465 |
| Mean respiratory PLVs (Respiratory condition) – Mean General Musical Sophistication scores |  | Pearson's r |  | | -0.121 |
|  |  | p-value |  | | 0.517 |

| **SI. Table 15 - Correlation between mean tone onset asynchronies and mean General Musical Sophistication scores** | | | | | |
| --- | --- | --- | --- | --- | --- |
| Variables |  | | |  | |
| Mean onset asynchronies (Normal condition) – Mean General Musical Sophistication scores |  | Pearson's r |  | | -0.024 |
|  |  | p-value |  | | 0.896 |
| Mean onset asynchronies (Auditory condition) – Mean General Musical Sophistication scores |  | Pearson's r |  | | 0.091 |
|  |  | p-value |  | | 0.625 |
| Mean onset asynchronies (Respiratory condition) – Mean General Musical Sophistication scores |  | Pearson's r |  | | -0.008 |
|  |  | p-value |  | | 0.967 |

| **SI, Table 16 - Correlation between mean cardiac PLVs and mean years of private instrument training** | | | | | |
| --- | --- | --- | --- | --- | --- |
| Variables |  | | |  | |
| Mean cardiac PLVs (Normal condition) – Mean years of private instrument training |  | Pearson's r |  | | 0.053 |
|  |  | p-value |  | | 0.776 |
| Mean cardiac PLVs (Auditory condition) – Mean years of private instrument training |  | Pearson's r |  | | 0.110 |
|  |  | p-value |  | | 0.556 |
| Mean cardiac PLVs (Respiratory condition) – Mean years of private instrument training |  | Pearson's r |  | | -0.119 |
|  |  | p-value |  | | 0.523 |

| **SI. Table 17 - Correlation between mean respiratory PLVs and mean years of private instrument training** | | | | | |
| --- | --- | --- | --- | --- | --- |
| Variables |  | | |  | |
| Mean respiratory PLVs (Normal condition) – Mean years of private instrument training |  | Pearson's r |  | | -0.162 |
|  |  | p-value |  | | 0.384 |
| Mean respiratory PLVs (Auditory condition) – Mean years of private instrument training |  | Pearson's r |  | | -0.041 |
|  |  | p-value |  | | 0.825 |
| Mean respiratory PLVs (Respiratory condition) – Mean years of private instrument training |  | Pearson's r |  | | -0.156 |
|  |  | p-value |  | | 0.402 |

| **SI. Table 18 - Correlation between mean tone onset asynchronies and mean years of private instrument training** | | | | | |
| --- | --- | --- | --- | --- | --- |
| Variables |  | | |  | |
| Mean tone onset asynchronies (Normal condition) – Mean years of private instrument training |  | Pearson's r |  | | -0.031 |
|  |  | p-value |  | | 0.869 |
| Mean tone onset asynchronies (Auditory condition) – Mean years of private instrument training |  | Pearson's r |  | | -0.044 |
|  |  | p-value |  | | 0.816 |
| Mean tone onset asynchronies (Respiratory condition) – Mean years of private instrument training |  | Pearson's r |  | | -0.175 |
|  |  | p-value |  | | 0.346 |

**SI Part II: Social Interaction Questions**

**13. Social interaction ratings**

We assessed participants' subjective perceptions of interpersonal coordination by evaluating four questions collected after each Production condition: pleasantness, connectedness, relationship, and synchronization success on a 7-point Likert scale (1 = lowest pleasantness/connectedness/relationship/success, 7 = highest pleasantness/connectedness/relationship/success). Separate ANOVAs addressed each question in terms of the 3 conditions (Auditory, Respiratory, and the averaged Normal conditions).

Significant differences were found in participants' ratings of pleasantness (*F* (2, 60) = 6.272, *p* = 0.003, *η²* = 0.173) and of synchronization success (*F* (2, 60) = 11.139, *p* < .001, *η²* = 0.271). No significant differences were observed in connectedness or relationship (SI Table 21, 22). Pleasantness ratings that followed the Respiratory condition showed significantly lower scores (mean = 4.855) than ratings that followed the Auditory condition (mean = 5.339, *t* (30) = 2.752, *p_holm_* = 0.020, *d* = 0.584) or the Normal condition (mean = 5.274, *t*(30) = 2.94, *p_holm_* = 0.019, *d* = 0.506). Similarly, synchronization success ratings that followed the Respiratory condition were significantly lower (mean = 4.435) than ratings that followed the Auditory condition (mean = 5.081, t(30) = 3.616, *p_holm_* = 0.002, *d* = 0.672) or the Normal condition (mean = 5.169, *t*(30) = 4.680, *p_holm_* < .001, *d* = 0.975). No significant differences were observed between the Auditory and Normal conditions in pleasantness or synchronization success scales. Consistent with the behavioral synchronization results, these findings suggest that the Respiratory condition posed the greatest challenge for partners.

***SI Table 19 - Q1 (Pleasantness):**

| *Within Subjects Effects* | | | | | | | | | | | | |
| --- | --- | --- | --- | --- | --- | --- | --- | --- | --- | --- | --- | --- |
| Cases | | Sum of Squares | | df | | Mean Square | | F | | p | | η² |
| Condition |  | 4.280 |  | 2 |  | 2.140 |  | 6.272 |  | 0.003 |  | 0.173 |
| Residuals |  | 20.470 |  | 60 |  | 0.341 |  |  |  |  |  |  |
| Note.  Type III Sum of Squares | | | | | | | | | | | | |

| *Post Hoc Comparisons - Condition* | | | | | | | | | | | | | | |
| --- | --- | --- | --- | --- | --- | --- | --- | --- | --- | --- | --- | --- | --- | --- |
|  | |  | | Mean Difference | | SE | | df | | t | | Cohen's d | | p_holm_ |
| Normal |  | Auditory |  | -0.065 |  | 0.122 |  | 30 |  | -0.531 |  | -0.078 |  | 0.600 |
|  |  | Respiratory |  | 0.419 |  | 0.143 |  | 30 |  | 2.940 |  | 0.506 |  | 0.019 |
| Auditory |  | Respiratory |  | 0.484 |  | 0.176 |  | 30 |  | 2.752 |  | 0.584 |  | 0.020 |
| Note.  P-value adjusted for comparing a family of 3 estimates. | | | | | | | | | | | | | | |

**SI Table 20 - Q4 (Synchronization Success):**

| *Within Subjects Effects* | | | | | | | | | | | | |
| --- | --- | --- | --- | --- | --- | --- | --- | --- | --- | --- | --- | --- |
| Cases | | Sum of Squares | | df | | Mean Square | | F | | p | | η² |
| Condition |  | 14.215 |  | 2 |  | 7.108 |  | 11.139 |  | < .001 |  | 0.271 |
| Residuals |  | 38.285 |  | 60 |  | 0.638 |  |  |  |  |  |  |
| Note.  Type III Sum of Squares | | | | | | | | | | | | |

| *Post Hoc Comparisons - Condition* | | | | | | | | | | | | | | |
| --- | --- | --- | --- | --- | --- | --- | --- | --- | --- | --- | --- | --- | --- | --- |
|  | |  | | Mean Difference | | SE | | df | | t | | Cohen's d | | p_holm_ |
| Normal |  | Auditory |  | 0.290 |  | 0.211 |  | 30 |  | 1.376 |  | 0.302 |  | 0.179 |
|  |  | Respiratory |  | 0.935 |  | 0.217 |  | 30 |  | 4.307 |  | 0.975 |  | < .001 |
| Auditory |  | Respiratory |  | 0.645 |  | 0.178 |  | 30 |  | 3.616 |  | 0.672 |  | 0.002 |
| Note.  P-value adjusted for comparing a family of 3 estimates. | | | | | | | | | | | | | | |

**SI Table 21 - Q2 (Connectedness):**

| *Within Subjects Effects* | | | | | | | | | | | | |
| --- | --- | --- | --- | --- | --- | --- | --- | --- | --- | --- | --- | --- |
| Cases | | Sum of Squares | | df | | Mean Square | | F | | p | | η² |
| Condition |  | 0.808 |  | 2 |  | 0.404 |  | 1.493 |  | 0.233 |  | 0.047 |
| Residuals |  | 16.234 |  | 60 |  | 0.271 |  |  |  |  |  |  |
| Note.  Type III Sum of Squares | | | | | | | | | | | | |

| *Post Hoc Comparisons -* Condition | | | | | | | | | | | | | | |
| --- | --- | --- | --- | --- | --- | --- | --- | --- | --- | --- | --- | --- | --- | --- |
|  | |  | | Mean Difference | | SE | | df | | t | | Cohen's d | | p_holm_ |
| Normal |  | Auditory |  | -0.202 |  | 0.143 |  | 30 |  | -1.409 |  | -0.248 |  | 0.508 |
|  |  | Respiratory |  | -0.008 |  | 0.105 |  | 30 |  | -0.077 |  | -0.010 |  | 0.939 |
| Auditory |  | Respiratory |  | 0.194 |  | 0.144 |  | 30 |  | 1.342 |  | 0.238 |  | 0.508 |
| Note.  P-value adjusted for comparing a family of 3 estimates. | | | | | | | | | | | | | | |

**SI Table 22- Q3 (Relationship):**

| *Within Subjects Effects* | | | | | | | | | | | | |
| --- | --- | --- | --- | --- | --- | --- | --- | --- | --- | --- | --- | --- |
| Cases | | Sum of Squares | | df | | Mean Square | | F | | p | | η² |
| Condition |  | 1.055 |  | 2 |  | 0.528 |  | 1.473 |  | 0.237 |  | 0.047 |
| Residuals |  | 21.497 |  | 60 |  | 0.358 |  |  |  |  |  |  |
| Note.  Type III Sum of Squares | | | | | | | | | | | | |

| *Post Hoc Comparisons - Condition* | | | | | | | | | | | | | | |
| --- | --- | --- | --- | --- | --- | --- | --- | --- | --- | --- | --- | --- | --- | --- |
|  | |  | | Mean Difference | | SE | | df | | t | | Cohen's d | | p_holm_ |
| Normal |  | Auditory |  | -0.036 |  | 0.122 |  | 30 |  | -0.298 |  | -0.030 |  | 0.768 |
|  |  | Respiratory |  | 0.206 |  | 0.142 |  | 30 |  | 1.448 |  | 0.172 |  | 0.474 |
| Auditory |  | Respiratory |  | 0.242 |  | 0.185 |  | 30 |  | 1.306 |  | 0.202 |  | 0.474 |
| Note.  P-value adjusted for comparing a family of 3 estimates. | | | | | | | | | | | | | | |

**Questionnaires**

**The questions below concern your impression of the experiment you just completed. Please answer all questions by circling a number on each of the scales. There are no right or wrong answers to any of these questions, we are just interested in your opinions.**

1. How unpleasant or pleasant did you find the experience of playing with the other participant?

|  | 1 | 2 | 3 | | 4 | 5 | 6 | | 7 |
| --- | --- | --- | --- | --- | --- | --- | --- | --- | --- |
| Extremely unpleasant | | | |  | | | | Extremely pleasant | |

2. Overall, I feel connected to the person in the experiment with me.

|  | 1 | 2 | 3 | | 4 | 5 | 6 | | 7 |
| --- | --- | --- | --- | --- | --- | --- | --- | --- | --- |
| Disagree Strongly | | | |  | | | | Agree Strongly | |

3. Please circle the picture below that best describes how your relationship with the other person in the experiment currently is:

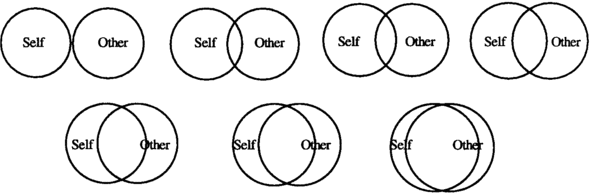

4. How successful do you think your synchronization was with your partner?

|  | 1 | 2 | 3 | | 4 | 5 | 6 | | 7 |
| --- | --- | --- | --- | --- | --- | --- | --- | --- | --- |
| Not at all  synchronized | | | |  | | | | Very well synchronized | |
